# A molecular gradient along the longitudinal axis of the human hippocampus informs large-scale behavioral systems

**DOI:** 10.1101/587071

**Authors:** Jacob W. Vogel, Renaud La Joie, Michel J. Grothe, Alex Diaz-Papkovich, Andrew Doyle, Etienne Vachon-Presseau, Claude Lepage, Reinder Vos de Wael, Yasser Iturria-Medina, Boris Bernhardt, Gil D. Rabinovici, Alan C. Evans

**Affiliations:** Montreal Neurological Institute, McGill University, Montŕeal, QC, Canada; Memory and Aging Center, University of California, San Francisco, CA, USA; German Center for Neurodegenerative Diseases (DZNE), Rostock, Germany; McGill University, Montŕeal, QC, Canada

**Keywords:** Hippocampus, Gene expression, Neuroimaging, Brain Networks, Imaging Genetics, Neurodegenerative Disease, Cognition

## Abstract

The functional organization of the hippocampus is distributed as a gradient along its longitudinal axis that explains its differential interaction with diverse brain systems. We show that the location of human tissue samples extracted along the longitudinal axis of the hippocampus can be predicted within 2mm using the expression pattern of less than 100 genes. When variation in this specific gene expression pattern was observed across the whole brain, a distinct anterioventral-posteriodorsal gradient was observed. Frontal, anterior temporal and brainstem regions involved in social and motivational behaviors, selectively vulnerable to frontotemporal dementia and more functionally connected to the anterior hippocampus could be clearly differentiated from posterior parieto-occipital and cerebellar regions involved in spatial cognition, selectively vulnerable to Alzheimers disease, and more functionally connected to the posterior hippocampus. These findings place the human hippocampus at the interface of two major brain systems defined by a single distinct molecular gradient. (148/150)

## 1. Introduction

A phylogenetically conserved and well connected structure involved in a diverse multitude of behaviors, the hippocampus provides an excellent base for studying the evolution of cognition. Alongside its highly nuanced and well documented role in memory, the hippocampus has been implicated in many other behaviors and functions, ranging from social cognition to spatial orientation to regulation of endocrine processes, such as stress response [5, 8]. The hippocampus can be divided into well-described subfields – the cornu ammoni (CA), dentate gyrus and subiculum – which represent its principal axis of organization, and which strongly inform cytoarchitectonic variation and both internal and external circuitry [5]. A second orthogonal axis of organization of the hippocampus lies along its longitudinal axis in a gradient spanning its two poles. In the rodent, this axis is often referred to as the ventral-dorsal axis, while a homologous gradient is thought to exist in humans along the anterior-posterior axis [49, 20, 42]. To study variations along this axis, the hippocampus is often divided into basic macroscopic partitions; the head-body-tail division is often used in humans, whereas a dorsal-ventral division is used in rodents. The divisions along the longitudinal axis of the hippocampus are characterized by a complex but distinct pattern of afferent and efferent connections, as well as impressive behavioral domain specificity. In rodents, the ventral hippocampus shares connections with the prefrontal cortex, basolateral amygdala, hypothalamus, and other structures mediating neuroendocrine and autonomic signaling and motivated behavior. Meanwhile, the dorsal hippocampus is anatomically connected with retrosplenial cortex, mamillary bodies, anterior thalamic complex and other networks implicated in movement, navigation and exploration ([8, 20]). Studies directly assessing the existence of a homologous longitudinal organizational axis in the human hippocampus have found compelling evidence in support [52, 10, 14, 1], and evidence has emerged suggesting this axis defines the multifaceted role of the hippocampus in complex cognitive systems [44] and in vulernability to neurodegenerative diseases [28, 33].

Centrally involved in so many aspects of brain function and dysfunction, a comprehensive study of the hippocampus and its organizational principles may be paramount to understanding the brain at large. With this concept in mind, several studies have explored the molecular properties regulating the longitudinal axis of the hippocampus. A number of studies have characterized the genomic anatomy of the ventral-dorsal axis of the rodent hippocampus as a whole or across specific subfields [13, 15, 19, 31, 51], how gene expression along the axis changes over the course of development [29, 46], and how it influences patterns of connectivity [8]. While some consensus over implicated genes has been met, all of these studies have been performed exclusively in rodents, and it is unclear whether similar genes and proteins are responsible for regulating and characterizing the anterior-posterior axis of the human hippocampus. This distinction is important, as the human hippocampus bears a different anatomy from that of rodents, participates in ostensibly more complicated cognitive systems, and shows selective vulnerability to diseases unique to humans.

As yet, such explorations have been severely limited due to the complications of measuring regionally detailed gene expression in the human brain. However, the Allen Human Brain Atlas has provided unprecedented access to human brain gene expression data. In the current study we leverage gene expression data from the Allen Human Brain Atlas dataset to define the genomic anatomy of the longitudinal axis of the human hippocampus. Specifically, we sought to understand whether, as with the rodent hippocampus, notable gene expression variations also exists along the human hippocampus, and which genes are most prominently involved in this molecular organization. We further aimed to understand whether information about gene expression can help explain interactions between the hippocampus and the diverse brain systems it is associated with, as well as differential vulnerability to neurodegenerative disease. To accomplish this, we drew from several public and private human datasets to bridge molecular properties with brain structure and function, behavior, and finally, dissociated vulnerability to neurodegenerative disease. We show that a graduated pattern of gene expression along the hippocampal longitudinal axis predicts the location of a brain tissue sample along this axis, and that distinct interactions between the anterior and posterior hippocampus with specific brain systems can be predicted by the genomic similarity shared between those brain systems and the different poles of the hippocampus.

## 2. Results

### 2.1. A sparse set of genes can predict sample location along the longitudinal axis of the hippocampus

Normalized gene expression information from 58,692 probes were obtained from each of 170 brain samples extracted from the hippocampi of six deceased human donors from the Allen Human Brain Atlas. The longitudinal axis of the hippocampus, from the anterior to the posterior pole, was defined as a curve passing through the center of mass of the hippocampal volume of an average brain template in MNI standard space. The position of each of the 170 hippocampus samples was projected onto this longitudinal axis **(Fig. 1A, S1B)**. LASSO-PCR was used to create a model predicting the position of each sample based on its gene expression profile **(Fig. S1)**.

**Figure 1:**
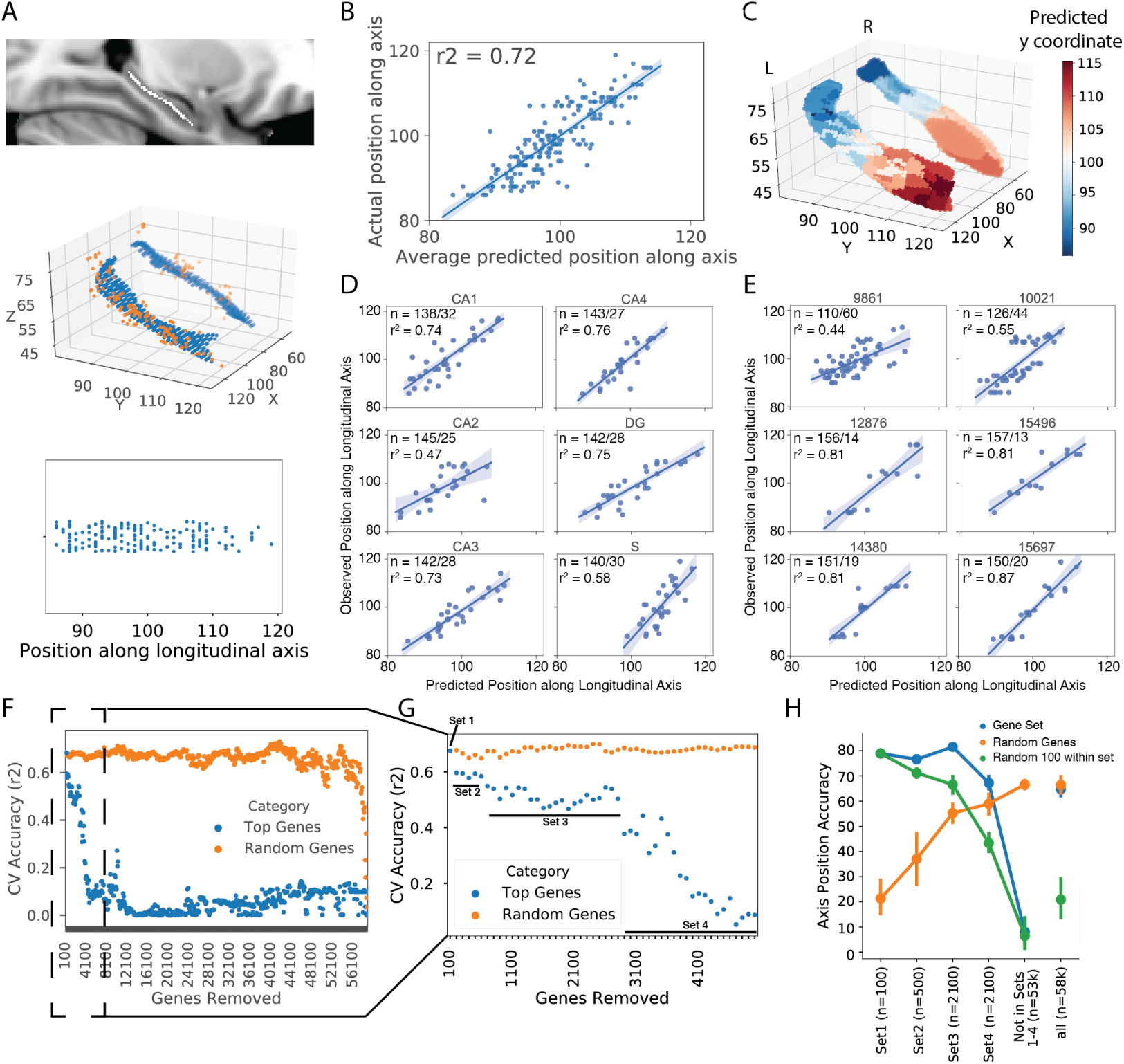
Gene expression predicts the location of tissue samples along the longitudinal axis of the hippocampus. A) (top) A curved skeleton of voxels was fitted along the center of mass of the hippocampal volume. (middle) Tissue samples (orange) were matched to the closest skeleton voxels (blue). (bottom) A sample’s position along the longitudinal axis was represented as the y-axis coordinate of the sample’s matched skeleton-voxel. B) Average predicted sample position (using gene expression) across ten separate 10-fold cross-validated LASSO-PCR models, compared to the actual position. C) Render of the hippocampal surface where each vertex shows the predicted location of the closest (surface projected) sample to that vertex. The smooth appearance of the right hippocampus is related to the fact that less samples were available for this structure. (D) Predicted vs. observed sample locations for leave-one-subfield-out models. For example, subpanel “CA1” shows the predicted vs. observed position of samples extracted from CA1 (test set) when the model was trained without CA1 samples (training set). In each plot, N represents the number of samples in the training and test sets. E) Predicted vs. observed sample locations for leave-one-donor-out models. F) The 100 most important probes in the LASSO-PCR model were iteratively removed and, after each removal, 10-fold cross-validation accuracy predicting sample position along the longitudinal axis was recorded (blue dots). G) The first 50 rounds of 100-probe removal from Panel A. Inflection points were identified after removing 100, 600, and 2700 genes. H) Accuracy in predicting sample position was recorded for models using different gene sets identified by the inflection points in panel G (blue), samples of 100 random within-set probes (green), and samples of random probes (orange) as input. Each model was run ten times with different bootstrap samples to calculate confidence intervals.

Using repeat ten fold cross-validation, the LASSO-PCR model explained 68-73% of the variance in sample position along the longitudinal axis (average MAE = 2.17mm) using only gene expression information **(Fig. 1B,C)**. The explained variance rose to 89% when the model was fit across all data.

By training our model on five subfields and then using this model to predict the position of the sixth left-out subfield (i.e. leave-one-subfield-out), we revealed that the genomic signature underlying the anterior-posterior gradient of the hippocampus is consistent across hippocampal subfields **(Fig. 1D)**, though the variance predicted was poorer for CA2 (r^2^ = 0.47) and the subiculum (r^2^ = 0.58) compared to CA1, CA3, CA4 and the dentate gyrus (r^2^s > 0.73). Leave-one-donor-out prediction additionally suggested consistency of the genomic signature across individuals **(Fig. 1E)**: while two donors accounted for over 60% of the samples, when samples from these two donors were included in the model, prediction of the location of samples for the other four donors was highly accurate (r^2^s > 0.80).

Weights from the LASSO-PCR model were back-transformed onto the individual probes in order to highlight the contribution of individual genes to the regulation of the hippocampal longitudinal axis. Weights from L1-regularized regression (LASSO) are difficult to reliably interpret [25], making identification of individual candidate genes challenging. To circumvent this issue, we iteratively removed the probes with 50 highest (anterior-associated) and 50 lowest (posterior-associated) weights, respectively, refit the model, and measured cross-validation accuracy of the new model, until all 58,692 probes were removed **(Fig. 1F)**. Removing the first set of 100 probes (Set 1) resulted in a sharp drop in cross-validation accuracy that was never recovered, supporting the notion that this gene set is important for regulating the longitudinal axis of the human hippocampus. Accuracy dropped once again after removing the next 500 probes (Set 2; rank 101-600), and after the next 1100 probes were removed (Set 3; rank 601-2700), cross-validation accuracy began to drop precipitously, finally bottoming out after another 2100 probes (Set 4; rank 2700-4800) were removed **(Fig. 1F,G)**. In contrast, iteratively removing sets of 100 random probes resulted in a very gradual and sporadic decrease in accuracy that only bottomed out when nearly all probes were removed **(Fig. 1F)**. Refitting the LASSO-PCR model with only probes from Set 1 (100 probes), Set 2 (500 probes) or Set 3 (2100 probes) resulted in cross-validation accuracy above 80% (MAE: Set1 = 1.84 mm; Set2 = 2.39 mm; Set 3 = 1.85 mm), a substantial improvement over the original model and a considerable improvement over models with equal-sized sets of random genes. Genes from Set 4 (2100 probes) alone achieved accuracy similar to a model using all (58,692) probes, and a model using all 53,892 probes not included in Sets 1-4 achieved cross-validation accuracy near 0% **(Fig. 1H)**. These results indicate that 100 specific probes are sufficient to accurately predict the location of a sample along the longitudinal axis of the hippocampus, and that probes outside of a specific set of 4800 provide little to no information about the axis. Fitting the model using gene Sets 2 and 3 alone resulted in cross-validation accuracy similar to Set 1, suggesting the possibility that important regulatory genes may also be present within these probe sets. However, the accuracy may also be assisted by the larger number of probes included in these two sets. Indeed, random sets of 100 probes taken from within Sets 2 and 3 showed reduced cross-validation accuracy compared to Set 1 and full Sets 2 and 3 **(Fig. 1H)**.

### 2.2. Candidate genomic regulators of the longitudinal axis of the human hippocampus

A list of the 100 top probes can be found in Table 1. Gene ontology (GO) enrichment analysis of the top 100 probes from the model (Set 1) revealed a consistent set of terms relating to regulation of anatomical structure morphogenesis and tissue (particularly axonal) growth and development. **(Fig. 2A)**. This gene set also included several genes previously identified to differentiate the dorsal and ventral aspects of the rodent hippocampus (e.g. NR2F2, SERTAD4, GDA, TTR, TPBG, SSTR1, TNNT2). Among this gene set, a feature explainer based on cross-validated Random Forest Regression suggested NR2F2 and RSPH9 as, on average, the most important local predictors of position along the longitudinal hippocampus axis **(Fig. 2C)**. This result remained consistent when additionally adding all probes from Sets 2 and 3 **(Supplementary Fig. S2)**. In addition to NR2F2 and RSPH9, the feature explainer also implicated local contributions to individual samples from FAM43B, FSTL4 and NTN1 **(Fig. 2C)**. The expression pattern of these five genes differed, as each pattern likely added unique information to the model **(Fig. 2D)**. For example, for some genes the anterior-posterior expression pattern was greater in certain subfields **(Fig. 2E)**.

**Table 1:** The top 50 anterior- and posterior-associated probes, respectively, identified by the LASSO-PCR model

| Anterior |  |  | Posterior |  |  |
| --- | --- | --- | --- | --- | --- |
| Probe | Gene | Beta | Probe | Gene | Beta |
| 1053204 | SERPINF1 | 0.0261 | 1023030 | NPNT | -0.0147 |
| 1053205 | SERPINF1 | 0.0253 | 1023031 | NPNT | -0.0141 |
| 1030761 | KLK7 | 0.0193 | 1050553 | TTR | -0.0129 |
| 1033144 | RSPH9 | 0.0170 | 1064147 | A_32.P11262 | -0.0129 |
| 1030762 | KLK7 | 0.0162 | 1058844 | BDKRB1 | -0.0124 |
| 1036045 | LYPD1 | 0.0145 | 1041274 | SERTAD4 | -0.0123 |
| 1041466 | GABRQ | 0.0144 | 1048608 | NTN1 | -0.0122 |
| 1032692 | PYDC1 | 0.0143 | 1039873 | HHIP | -0.0119 |
| 1042620 | SYTL2 | 0.0139 | 1039872 | HHIP | -0.0111 |
| 1034086 | RP13-102H20.1 | 0.0139 | 1017013 | RP11-561O23.6 | -0.0110 |
| 1042619 | SYTL2 | 0.0137 | 1038748 | GRHL2 | -0.0107 |
| 1051105 | SSTR1 | 0.0135 | 1041038 | RGMA | -0.0107 |
| 1041090 | LXN | 0.0134 | 1058843 | BDKRB1 | -0.0106 |
| 1031172 | TMEM215 | 0.0133 | 1042684 | BNC2 | -0.0105 |
| 1042621 | SYTL2 | 0.0133 | 1050668 | TPBG | -0.0104 |
| 1028032 | C1QL1 | 0.0132 | 1029814 | OSBPL3 | -0.0101 |
| 1010361 | PIRT | 0.0132 | 1048607 | NTN1 | -0.0100 |
| 1054831 | KCNG1 | 0.0132 | 1048537 | ONECUT2 | -0.0100 |
| 1059122 | AQP3 | 0.0130 | 1058080 | COL5A2 | -0.0100 |
| 1064467 | A_23.P213527 | 0.0128 | 1010982 | RP11-291L15.2 | -0.0099 |
| 1029570 | RP11-45B20.3 | 0.0128 | 1027004 | FSTL4 | -0.0098 |
| 1066217 | C1orf187 | 0.0126 | 1015986 | C1orf133 | -0.0098 |
| 1056223 | GPR39 | 0.0123 | 1048913 | DGKI | -0.0096 |
| 1021758 | OPRK1 | 0.0120 | 1010774 | DDC | -0.0096 |
| 1017426 | CD36 | 0.0119 | 1069644 | A_24.P401842 | -0.0096 |
| 1059123 | AQP3 | 0.0117 | 1070261 | A_32.P121537 | -0.0095 |
| 1030763 | KLK7 | 0.0117 | 1025477 | TNNT2 | -0.0095 |
| 1053962 | MYB | 0.0117 | 1027005 | FSTL4 | -0.0095 |
| 1056238 | GPR26 | 0.0116 | 1050554 | TTR | -0.0094 |
| 1054547 | LMO1 | 0.0115 | 1040196 | HPSE2 | -0.0094 |
| 1042988 | GPR88 | 0.0114 | 1012040 | DDC | -0.0093 |
| 1031384 | VGLL3 | 0.0114 | 1010523 | DDC | -0.0091 |
| 1014826 | NR2F2 | 0.0113 | 1058079 | COL5A2 | -0.0091 |
| 1013661 | NR2F2 | 0.0112 | 1038515 | WNT10A | -0.0090 |
| 1020068 | NR2F2 | 0.0112 | 1058569 | CASR | -0.0090 |
| 1046866 | GPR83 | 0.0111 | 1012029 | DDC | -0.0090 |
| 1048357 | GDA | 0.0110 | 1052410 | PVALB | -0.0089 |
| 1030949 | NRG1 | 0.0109 | 1060554 | A_24.P62668 | -0.0089 |
| 1031962 | RSPO2 | 0.0109 | 1033886 | FAM43B | -0.0088 |
| 1063851 | A_32.P136776 | 0.0108 | 1016934 | CTXN3 | -0.0088 |
| 1045386 | C20orf103 | 0.0108 | 1010582 | DDC | -0.0088 |
| 1037183 | SYTL1 | 0.0108 | 1040195 | HPSE2 | -0.0088 |
| 1054593 | LGALS2 | 0.0107 | 1043786 | GAL | -0.0087 |
| 1041091 | LXN | 0.0107 | 1039883 | GREM2 | -0.0087 |
| 1056237 | GPR26 | 0.0107 | 1026202 | KDELR3 | -0.0087 |
| 1013797 | KIAA1772 | 0.0107 | 1058081 | COL5A2 | -0.0086 |
| 1066971 | A_32.P115840 | 0.0106 | 1030360 | PDLIM5 | -0.0085 |
| 1048356 | GDA | 0.0106 | 1048538 | ONECUT2 | -0.0084 |
| 1033037 | SEMA3D | 0.0104 | 1060274 | A_24.P102119 | -0.0084 |
| 1020049 | NRG1 | 0.0104 | 1043787 | GAL | -0.0084 |

**Figure 2:**
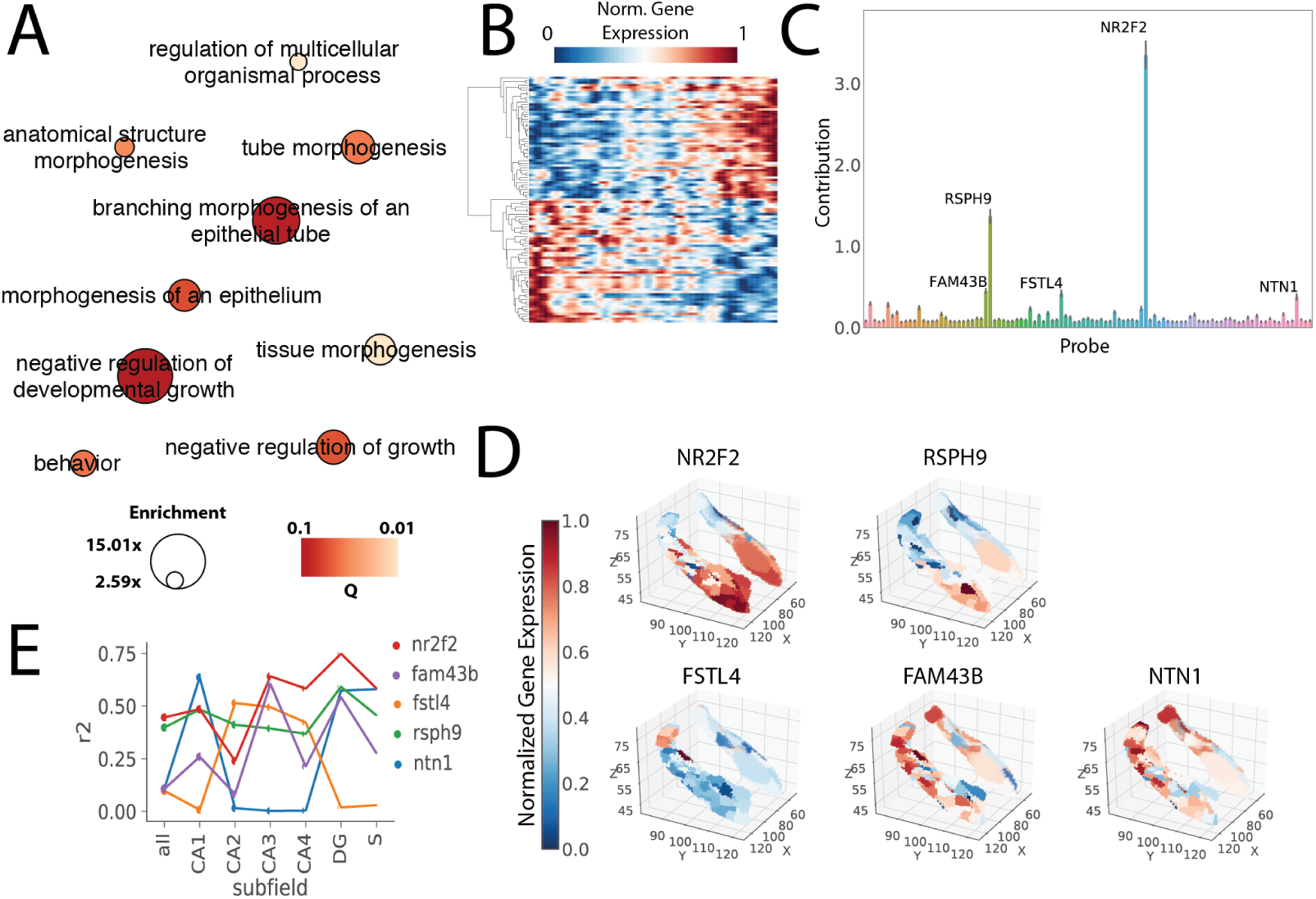
Candidate genes regulating the longitudinal axis of the human hippocampus. **A** Enriched Gene Ontology terms (Q<0.1) associated with Gene Set 1. Circle size indicates enrichment, whereas color indicates Q value (lighter = lower Q value). **B** Matrix showing gene expression for Gene Sets 1 (y-axis) across each hippocampal sample, ordered most posterior to most anterior (x-axis). Values were smoothed with a 3mm gaussian kernel across the x-dimension only and then clustered so that anterior-posterior patterns can be clearly visualized. **C** Average absolute local feature importances of probes in Gene Set 1 measured using a Random Forest-based feature explainer across all samples. **D** Surface rendering of the expression patterns of each of the five genes identified as locally important features to predicting position along the longitudinal axis. **E** For each of the five genes, the relationship between expression and position along the longitudinal axis (r^2^) is plotted stratified by subfield.

Feature explainers run on Sets 2 and 3 alone revealed more contributing features with less individual importance, compared to Set 1 and pools including Set 1 **(Supplementary Fig. S2)**. This suggests individual sample predictions are likely aided by different genes depending on their location along the longitudinal axis. Sets 2 and 3 may therefore contain a mix of genes regulating the longitudinal axis, genes regulated by the those genes, and genes that are independent but are specifically hyperexpressed in the anterior or posterior hippocampus. To partially explore this possibility, we performed GO enrichment analysis on all genes represented in Set 2, and then clustered genes sharing similar enrichment terms **(Supplementary Table S1)**. One cluster emerged sharing similar terms to those enriched in Set 1, relating to regulation of axon guidance, as well as cell motility, migration and development. This cluster also included genes previously described in studies exploring the rodent longitudinal axis, including SLIT2 and CADM1. Other GO enrichment sets included amine metabolic processes, GABA receptor activity, signal release/secretion, neuropeptide receptor activity, ion transport, behavior, serotonin receptor activity and lipoprotein mediated signaling. These latter gene clusters may be more likely to regulate behaviors differentially associated with the anterior or posterior hippocampus. We repeated this analysis for Set 3 **(Supplementary Table S2)**. Once again, a cluster of genes emerged associated with cell motility and migration, which again included genes previously described from the rodent literature (e.g. NTNG2, SEMA3E, NOV, SEMA4G, CADM1, CYP26B1). A second cluster emerged involving genes associated with both amine transport and neuronal migration, and also included some previously described genes (e.g. RAB3B, PENK, NTF3, NTS, OLFML2B, RASD2, RXRG, TIMP2).

As a way of validating the candidate genes identified, we repeated our analyses using Partial Least Squares regression (PLSR), another algorithm appropriate given the high dimensionality of our data. Using all probes, we obtained similar overall cross-validation results **(Supplementary Fig. S3)**. Of the top 100 probes identified by the PLSR model, 50 were included in Set 1, another 42 in Set 2, and the last 8 were found in Set3. Interestingly, of all probes in the model, NR2F2 and RSPH9 had the highest absolute beta estimates (weights), once again implicating these two genes as regulators of the longitudinal axis of the hippocampus **(Supplementary Table S3)**.

### 2.3. The genomic signature of the longitudinal axis of the hippocampus is represented as a spatial gradient across the brain

The Allen Human Brain Atlas data comprises 3702 samples across the brains of six donors. By leveraging the weights of our LASSO-PCR model, we created the Hippocampal Axis Genomic Gradient Index of Similarity (HAG-GIS), a value representing the degree to which the genomic signature of the hippocampal longitudinal axis is represented in the gene expression profile of a given non-hippocampus sample (**Fig. S1**). Larger positive values represent greater genomic similarity to the anterior hippocampus, while smaller negative values represent greater genomic similarity to the posterior hippocampus. When plotting these values for all brain samples, we observed a general pattern across the brain such that the brainstem and more antero-ventral sites of the cerebral cortex demonstrated greater genomic similarity to the anterior hippocampus, whereas the cerebellum and posterio-dorsal cortical regions demonstrated greater similarity to the posterior hippocampus **(Fig. 3, 4A)**.

**Figure 3:**
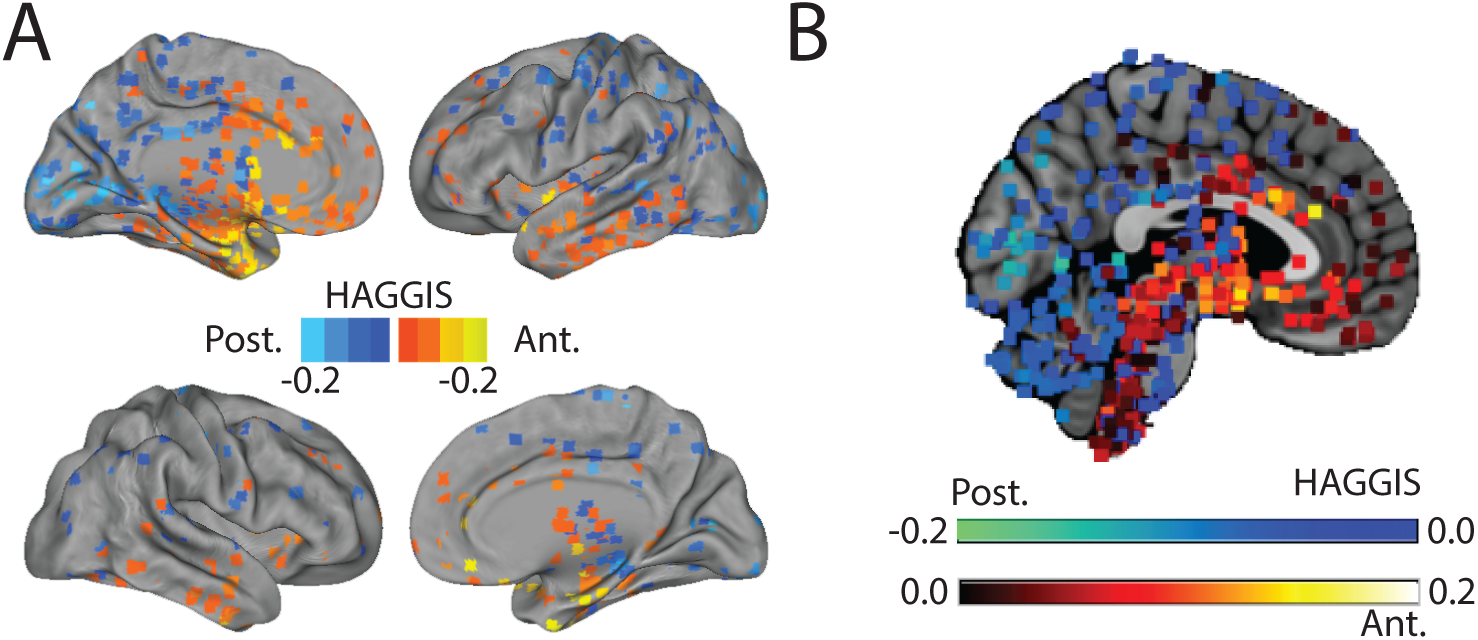
Spatial distribution of the HAGGIS across the brain. **(A)** Each sample was projected onto a cortical surface based on its MNI coordinates. Warm colors indicate the sample has a gene expression pattern more similar to the anterior hippocampus (higher HAGGIS), while cool colors represent the sample is more genomically similar to the posterior hippocampus (lower HAGGIS). **(B)** A medial slice inclusive of brainstem and cerebellum. Each dot represents a sample, and warm colors indicate higher HAGGIS, while cool colors represent lower HAGGIS. HAGGIS = Hippocampal Axis Genomic Similarity

**Figure 4:**
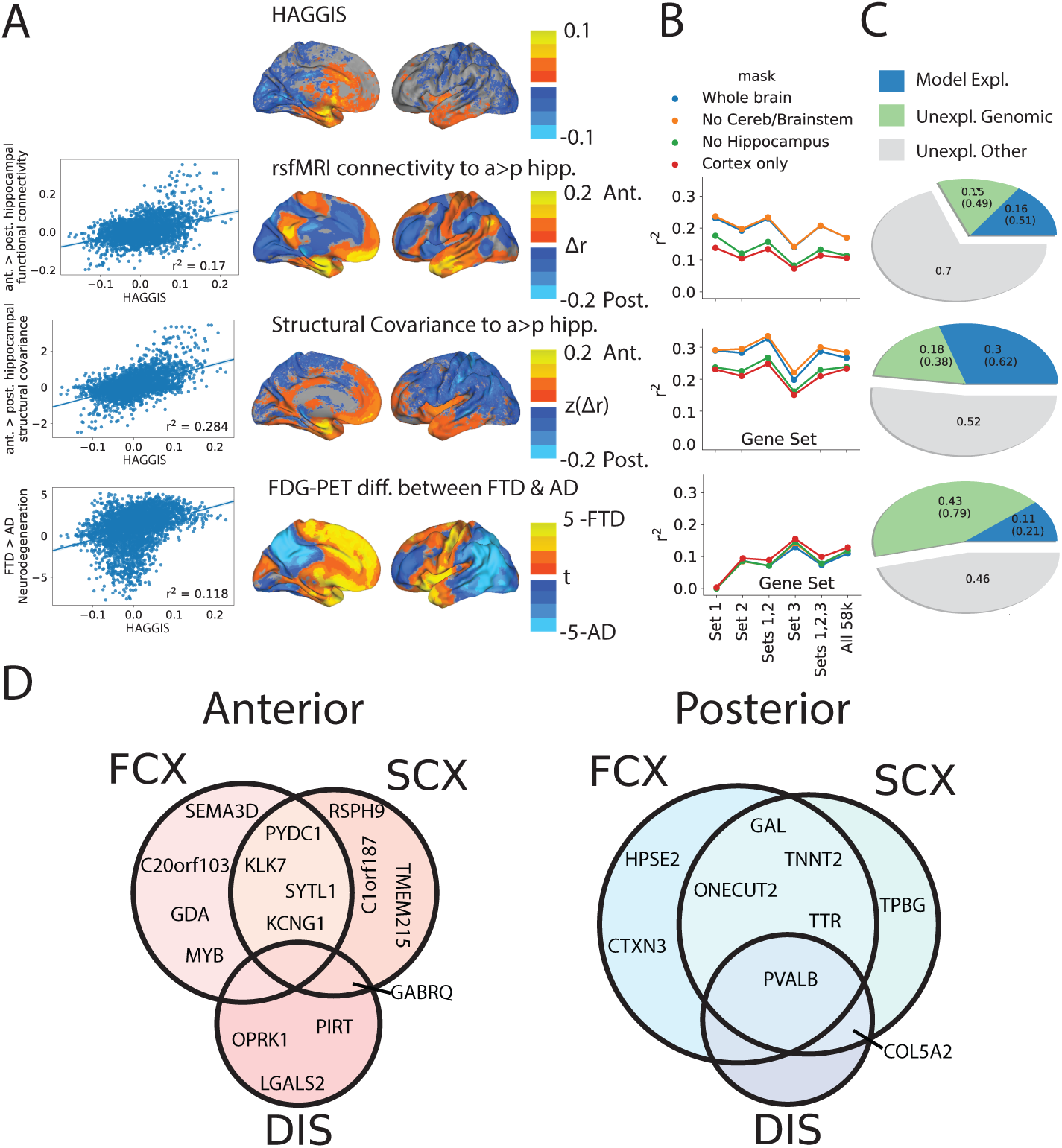
HAGGIS predicts hippocampus-brain relationships. **(A)** From top to bottom: The spatial distribution of (smoothed) HAGGIS across samples, differential functional connectivity to the anterior vs posterior hippocampus measured with rsfMRI (middle), differential structural covariance with the anterior vs posterior hippocampus, differential vulnerability to AD or FTD measured with FDG-PET. Graphs on the left visualize the relationship between these spatial patterns by comparing the HAGGIS of each sample with the mean value from the respective map within a 5-voxel cube around the sample coordinate. **B** Each of the above associations was re-calculated using three other brain masks, and using a HAGGIS formed from each gene set identified in Section 2.2. The r^2^ of each of these associations is visualized. **C** Pie charts indicating the proportion of genomic and total variance explained by each model. Numbers in parentheses indicate percentage of total genomic variance. **D** Genes involved in both the longitudinal axis of the hippocampus, and hippocampus-brain interactions. All genes pictured are among the top 50 anterior (red; top) or posterior (blue; bottom) features of the hippocampus longitudinal axis model. Each also participates in one or more hippocampus-brain interactions, indicated by the circles within the Venn diagrams. FCX = Differential functional connectivity between anterior and posterior hippocampus; SCX = Differential structural covariance between anterior and posterior hippocampus; DIS = Differential vulnerability between AD and FTD

### 2.4. Specific gene expression patterns inform interactions between the hippocampus and dissociated hippocampo-cortical systems

The anterior and posterior hippocampus each exhibit a distinct profile of anatomical connections in humans [1], which can also be represented using resting-state functional connectivity [52]. Using logistic regression and the HAGGIS, we identified coordinates to isolate the genomic posterior and anterior hippocampus **(Supplementary Fig. S4A)**. We then used an open database of resting-state functional connectivity information based on rsfMRI scans from 1000 subjects to create an average voxelwise map representing the degree to which brain regions are functionally connected to the anterior vs. posterior hippocampus. Brain samples bearing a gene expression profile more similar to the anterior hippocampus were also more functionally connected to this substructure, while the opposite pattern was observed for samples with gene expression profiles more similar to the posterior hippocampus (r^2^ = 0.170, **Fig. 4A**). A separate model was constructed in order to ascertain the maximum (cross-validated) variance in differential connectivity explainable given the (genomic) data. This analysis revealed that, while HAGGIS explained only 17% of the total model variance, it explained about 51% of the variance explainable with the present genomic data **(Fig. 4C**).

The strength of this relationship differed depending on where along the anterior-posterior axis the divisions were drawn, which parts of the brain were included, and the size of the cube used to extract data around the sample coordinate **(Supplementary Fig. S4C)**. The r^2^ ranged from 0.111 (central split, cortical only mask, 1mm cube diameter) to 0.304 (split at anterior/poster extremes, mask excluding only brainstem and cerebellum, 11mm cube diameter), though in all cases the relationship was observed to be significantly greater than chance (95% CI of chance r^2^ < 0.004 for all conditions; data not shown). The relationship between HAGGIS and functional connectivity also varied slightly depending on the gene Set used **(Fig. 4B**). Remarkably, prediction of functional connectivity by HAGGIS performed just as well when the HAGGIS was created using the smaller Sets, with the highest values achieved when only the top 100 probes were used.

A diverging pattern of structural covariance with the rest of the brain has also been observed across the longitudinal axis of the hippocampus [39], perhaps representing co-variation in cytoarchitecture. We used an open dataset of 153 structural MRI images from young healthy controls to create a map representing variation in structural covariance between the brain and the anterior vs posterior hippocampus. The more similar a brain region’s gene expression patterns were to the anterior hippocampus, the greater the structural covariance was between that structure and the anterior hippocampus, and vice versa for the posterior hippocampus (r^2^ = 0.284; **Fig. 4A**). HAG-GIS explained 62% of the variance explainable with the present genomic data **Fig. 4C**). This relationship varied but remained strong across different brain masks and gene sets **(Fig. 4B**).

To validate these finding without relying on an anterior-posterior split, we utilized a previously validated data-driven approach [52, 36] to extract the principal gradients of hippocampal functional connectivity and structural covariance with the rest of the brain, respectively. We then tested the relationship between each gradient and the predicted location of each sample based on the HAGGIS (**Supplementary Table S4**). For structural covariance, the 1st gradient, explaining 24% of the total variance in brain-hippocampus covariance, showed a strong correlation with HAGGIS (r^2^=0.41; **Supplementary Fig. S4D**). For functional connectivity, the 3rd gradient, explaining 13.5% of the total variance of hippocampus-brain connectivity, also showed a strong relationships with HAGGIS (r^2^=0.40; **Supplementary Fig. S4E**). These findings were not contingent on the gene set used to calculate the HAGGIS **(Supplementary Fig. S4F**).

### 2.5. Variation in genomic signature predicts regional vulnerability to neurodegenerative disease

The anterior and posterior hippocampus are also differentially involved in disparate neurodegenerative diseases [29], particularly Alzheimer’s disease (AD) and frontotemporal dementia (FTD) [44, 28, 33]. We acquired fluorodeoxyglucose (FDG) PET scans measuring glucose metabolism, a measure of neuronal health and degeneration, from patients diagnosed in a tertiary memory clinic as having AD or FTD. We used these scans to create a statistical map representing the relative patterns of neurodegeneration in AD vs. FTD. We found that samples with greater genomic similarity to the anterior hippocampus also showed greater hypometabolism in FTD compared to AD, whereas samples more similar to the posterior hippocampus showed greater hypometabolism in AD compared to FTD (r^2^ = 0.118; **Fig. 4A**). HAGGIS explained about 21% of the variance explainable given the present genomic information **(Fig. 4C**). This relationship also varied depending on the regions included and cube size, with r^2^ ranging from 0.095 (whole-brain, 1mm cube diameter) to 0.153 (cortex-only mask, 11mm cube diameter, **Supplementary Fig. S6)**, but remained greater than chance in all cases (data not shown). Notably, and unlike previous analyses, the relationship between HAGGIS and regional disease vulnerability was not observed when restricting the HAGGIS to the top 100 probes (Set 1) **(Fig. 4B)**.

### 2.6. Specific genes link longitudinal axis to connectivity and vulnerability patterns

In order to highlight specific genes that may be involved in both maintenance of the longitudinal axis and hippocampus-brain interaction, we constructed independent models to learn the genomic profile of the maps from **Fig. 4A** and compared the top 100 features from these models to the longitudinal axis model. The proportion of overlap between the top 100 features of each model with the top 100 features from the hippocampus longitudinal axis model far exceeded chance (Functional: 20%; Structural: 21%; Disease: 11%). Overlapping genes from each model, stratified by involvement in anterior or posterior hippocampus, can be found in **Fig. 4D**. Interestingly, some genes were involved in multiple systems. For example, PVALB, specifically expressed in the posterior hippocampus, was also highly expressed in brain regions functionally connected and structurally covarying with the posterior hippocampus, as well as in regions specifically vulnerable to Alzheimer’s disease. Additionally, anterior hippocampus gene GABRQ was also highly expressed in regions both structurally covarying with the anterior hippocampus and those vulnerable to frontotemporal dementia.

### 2.7. Variation in genomic signature predicts involvement in distributed cognitive networks

The posterior and anterior hippocampus are implicated in distinct aspects of memory and cognition [52, 10, 14, 16]. We explored whether regions sharing genomic similarity to the posterior or anterior hippocampus were more likely to participate in cognitive networks proposed to involve those substructures. We downloaded 100 meta-analytic functional coactivation maps from the Neurosynth database, each composed from between 91 and 4201 task-based functional MRI activation studies, and each of which was paired with a set of related cognitive/behavioral topics. These topic/map pairs represent greater-than-chance regional functional coactivation patterns reported consistently in studies sharing words from certain related topic-sets in the publication text. These maps can therefore be thought to represent specific region-sets involved in distributed cognitive networks. We calculated the mean HAGGIS of samples falling within each cognitive map, with higher positive values indicating greater genomic covariance between the regions covered by that coactivation map and the anterior hippocampus, and lower negative values representing greater genomic covariance with the posterior hippocampus.

Using a conservative approach (only including maps with at least 500 overlapping samples: 29 maps; minimum map size: 36,622 voxels), we observed a pattern largely consistent with previous hypotheses of hippocampal involvement in different cognitive systems [44] **(Fig. 5)**. As we hypothesized, regions that expressed a gene expression profile more consistent with the anterior hippocampus tended to be those involved in social and emotional cognition, but also included maps associated with reward and conditioning, among others. Also consistent with our hypotheses, cognitive networks more genomically similar to the posterior hippocampus were associated with spatial cognition, imagination and mental simulation, but also included maps associated with visualization, working memory and movement/action. Interestingly, maps associated with episodic memory and physical stimulation slightly favored the anterior hippocampus, or were not strongly associated with either posterior or anterior hippocampus. These patterns remained remarkably similar when repeating the analysis with only probes from Set 1, representing the top 100 probes in our model **(Supplementary Fig. S5)**.

**Figure 5:**
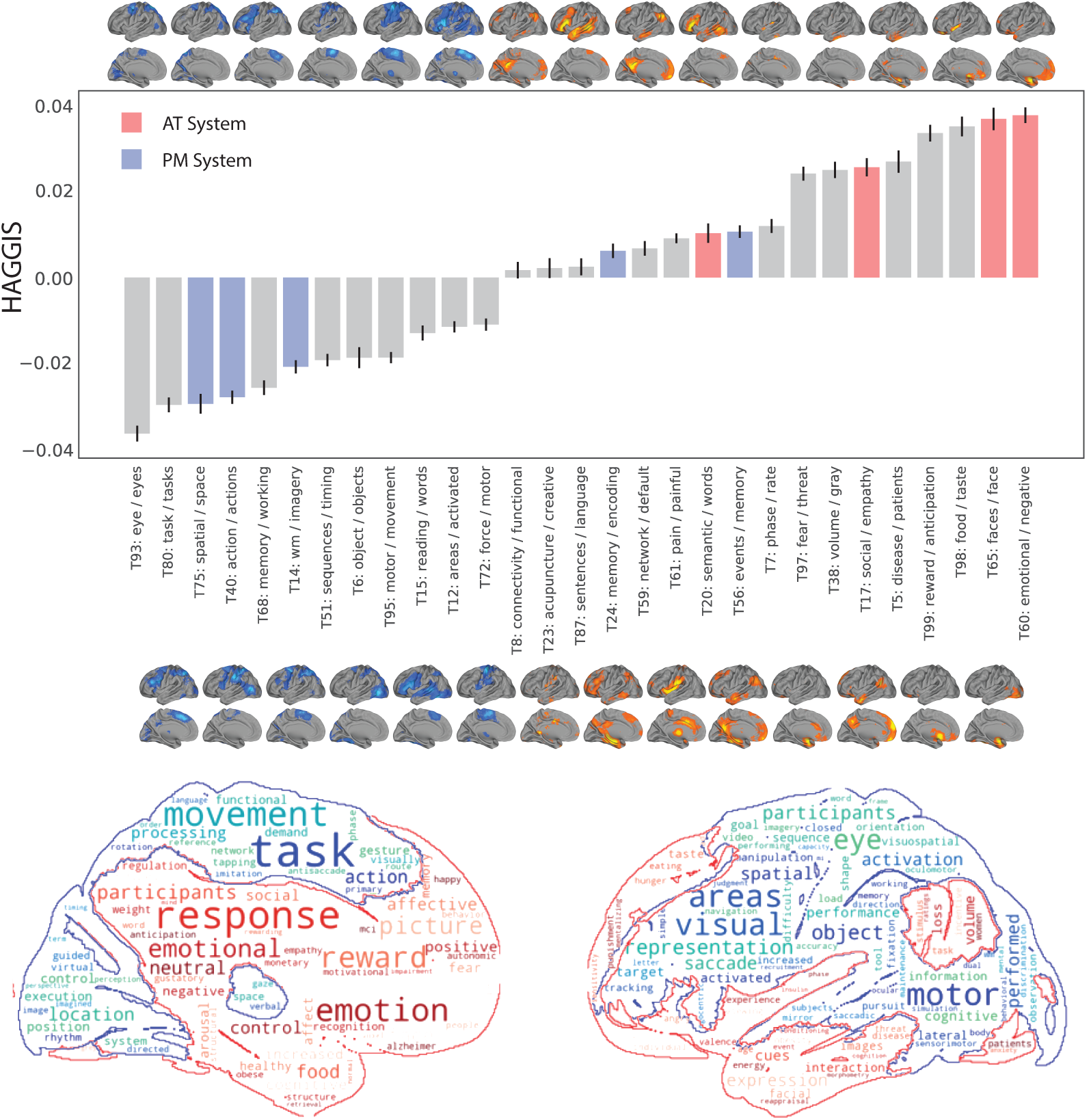
Variation in genomic signature predicts involvement in distributed cognitive networks. (Top) Maps were downloaded from neurosynth representing greater than chance meta-analytic functional activation in studies with different topic-sets mentioned in their abstract. Mean HAGGIS (represented by bars) was calculated for samples inside maps encompassing >500 samples (visualized either directly above or directly below each bar). Error bars represent SEM. Topic hypothesized to belong to the AT or PM system are shown in red and blue respectively. A word cloud summarizing the regions and topics most associated with the genomic signal of the anterior (red) and posterior (blue) hippocampus

## 3. Discussion

The hippocampus plays a central role in many systems that regulate behavioral processes across several species, and that are dysregulated in several human neuropsychiatric diseases. The contribution of the hippocampus to many of these systems is grossly organized across its longitudinal atlas. Characterizing the molecular properties of this axis may be vital to understanding how gene expression networks regulate macroscopic brain networks. We show that the anterior-posterior position of a tissue sample extracted from the human hippocampus can be predicted with remarkable accuracy using the expression pattern of only a handful of genes. Further, we find genomic representation of the anterior-posterior gradient projected across the entire brain, and that this representation partially explains relationships between the hippocampus and dissociated hippocampo-cortical systems. The anterior hippocampus shares genomic patterning with a system encompassing the medial prefrontal cortex, anterior temporal lobe and the brainstem. In general, these regions showed greater functional connectivity and structural covariance with the anterior than the posterior hippocampus, greater vulnerability to FTD than to AD, and more frequent involvement in cognitive tasks involving motivation/conditioning, social and emotional cognition and semantic knowledge. The posterior hippocampus, in contrast, shared a genomic pattern with the cerebellum, and occipital, parietal and motor and pre-motor cortex. These regions generally showed greater connectivity and structural covariance with the posterior than anterior hippocampus, more vulnerability to AD than FTD, and were more likely to participate in cognitive tasks involving spatial representation, visual processing, working memory and simulation. These results confirm and extend findings across species and sub-disciplines of neuroscience to suggest shared gene expression patterns underlying a well-described dissociation of anterior vs posterior hippocampal involvement in cognitive brain networks. Further, the findings support the existence of a specific axis of organization in the human brain, where an anterior-ventral - posterior-dorsal gradient explains regional involvement in diverse behaviors, underscored by a specific pattern of gene expression. These findings together form a template for studying how specific genes may regulate the development of dissociated hippocampal connectivity networks in humans and their involvement in specific behaviors and, potentially, specific diseases.

Our results support an existing concept of molecular gradients in the cerebral cortex [11, 36, 21]. The anterior-ventral - posterior-dorsal pattern observed in our data is reminiscent of a general anterior-posterior molecular gradient previously observed in the Allen Human Brain Atlas dataset [26, 21]. [21] reviews qualities of this gradient, including a pattern of neuronal organization where, as one moves caudally to rostrally, neuron and arbor size increase while neuronal number and density decrease. Perhaps related, a “dual origin” hypothesis has been proposed suggesting the cerebral cortex has developed radially from certain phylogenetically conserved limbic structures over the course of evolution. This hypothesis describes a ventral system emanating from the perirhinal and amygdalar cortex that is involved in semantic identification of a stimulus and motivated behavior, while a dorsal system has evolved from the hippocampus and parahippocampal cortex to coordinate spatial representation and coordinated action [23]. The hypothesis is supported by evidence from comparative cytoarchitectonics and connectivity patterns across species. The AT/PM hypothesis [44] provides yet another example of opposing cortical systems loosely following an anterior-posterior organization and determining patterns of brain organization. Each of these models originated from a different field of inquiry (gene expression, cortical evolution, memory network organization), but the models converge in many respects, and a microcosm of this shared framework seems to be represented along the longitudinal axis of the hippocampus – explicitly so in the AT/PM and dual origin models. Our results generally support the premise that the hippocampus participates in two distinct macroscopic networks characterized by distinct structural covariance, functional connectivity, behavioral domain specificity and disease vulnerability, and that participation in these networks can be predicted by position along the longitudinal axis. However, we take this framework one step further to suggest these two distinct networks are composed of one single gradient of gene expression that may be underlying their systemic distinctions.

While our initial model utilized the expression patterns of nearly 60,000 probes corresponding to over 20,000 genes, the model favored a much smaller profile of probes to describe the longitudinal axis of the human hippocampus. When isolating a small set of only 100 probes, we were able to successfully predict the location of samples along the longitudinal atlas with less than 2 mm error, as well as interactions between the hippocampus and the specific brain systems. The set was enriched with genes associated with, in particular, anatomical structure morphogenesis and cellular growth, suggesting genes within this set may be involved in coordinating and/or maintaining the anatomical variation of the hippocampus along its longitudinal axis. Whether these genes are also partially responsible for the functional variation along the axis remains unclear, though it is notable that similar expression patterns of these 100 genes can be observed in other brain regions that interact with the hippocampus. In particular, we identified several specific genes that appear to be involved in coordinating both the longitudinal axis of the hippocampus and one or more aspects of the hippocampus-associated distributed brain networks. A number of these genes (PVALB, GAL, ONE-CUT2, PIRT, TNNT2, RSPH9, COL5A2, CTXN3) have been reported in previous studies examining genes regulating functional network organization [47, 54]. The shared gene expression across disparate regions may be related to shared anatomical characteristics (e.g.[3, 45, 48]), and examining the contribution of genes across these different network properties (structure, function, disease vulnerability) may lead to a clearer picture of the role various proteins play in overall network organization.

Many of the genes identified in our study have also been described in previous studies characterizing the dorsal and ventral subdivisions and longitudinal gradients of the rodent hippocampus (e.g. [31, 51, 29, 46, 15]). This suggests a fair degree of homology between rodents and human in the distribution of proteins along the longitudinal axis of the hippocampus, and perhaps in the development and maintenance of the axis itself. However, many previously undocumented proteins were also identified, and replication and comparative studies will be required to disentangle whether these candidate genes are truly unique to humans or a result of small sample sizes and differing methodologies.

The most important genes identified in our model can be interpreted as the most central genes in the hippocampal gene expression network(s) most associated with position along the longitudinal axis of the hippocampus. We cannot infer which genes are causally related to axis formation and maintenance and, as weights from backward regression problems are notoriously hard to interpret [25], even identifying the most important among a set of genes is challenging. Being aware of these limitations, we identified NR2F2 (also called COUP-TFII) and RSPH9 to be particularly important in local prediction of sample location along the axis. This likely suggests that these two genes demonstrated the cleanest and most consistent linear gradient in expression across the longitudinal axis among those assessed (which can be visually appreciated by the surface plots of expression levels of these genes across the hippocampus in Fig. 2D). The pattern of expression we observed here mirror descriptions of other studies of NR2F2 expression in the rodent hippocampus [22], as well as more macroscopically in the human brain during development, particularly in the temporal lobes [4]. NR2F2 is also key in the determination of cell fate in numerous circumstances [32, 27], including that of interneurons expressing PVALB (parvalbumin) or SST (somatostatin), where NR2F2 promotes SST and represses PVALB [27]. These findings are highly consistent with the expression of PVALB (expressed posteriorly) and SSTR1 (expressed anteriorly with NR2F2) in our data (Table 1). For its part, RSPH9 is part of the structure of primary cilia, which can be found within ependymal cells lining the ventricles, as well as in the CA1 subfield of the hippocampus and adjacent choroid plexus [56]. There is evidence that these cilia can promote neurogenesis in the hippocampus through mediating expression of SHH (sonic hedgehog) [9], a protein implicated in anterior-posterior pattern formation, and identified as a possible protein of importance in our data by the presence of HHIP (hedgehog-interacting protein) among the top 100 anterior-posterior associated genes (Table 1). The expression patterns of RSPH9 in our data may signal the presence of a specific pattern of cilia, which may help regulate the longitudinal axis of the hippocampus through hedgehog signaling. Both NR2F2 and RSPH9 have been identified as role-players in human functional network organization [47, 54].

The longitudinal axis of the hippocampus provides many clues about the development and characterization of different behavioral systems, which may be particularly important when it comes to understanding diseases characterized by selective dysfunction of these systems. Understanding the molecular components that maintain vulnerable systems may go a long way in learning which components are responsible when the system begins to fail. Data from multiple studies support a specific role for the longitudinal axis of the hippocampus in AD and FTD [28, 33]. Our data support this notion, suggesting that regions more vulnerable to FTD than AD share a more similar molecular profile to the anterior than posterior hippocampus, and that the opposite pattern was observed for regions more vulnerable to AD than FTD. It is tempting to wonder whether the same genes that coordinate the development of systems also incidentally contribute to the degeneration of these systems over time, a possible example of antagonistic pleiotropy. For example, among the top 100 anterior-posterior associated genes identified in our model were several genes known to interact with amyloid-*β* protein, the primary pathologic hallmark of AD, many of them specifically associated with the posterior hippocampus in our data. For example, TTR has been shown to bind amyloid-*β* aggregates in a chaperone-like manner [12], and TTR mutations have been associated with hippocampal atrophy in aging humans [17]. NTN1 interacts with the amyloid precursor protein (APP) and has been described as a key regulator of amyloid-*β* production [43, 34]. Much less is known about the molecular properties of FTD, but it was interesting to see the KLK7 gene among the top anterior hippocampus-associated genes (Table 1), as the KLK7 protein and other kallikreins have been found to be reduced in the CSF of FTD patients [18]. Although little can be extrapolated from our data about the potentially dissociated role of specific proteins in AD and FTD, we provide evidence for distinct molecular properties that characterize the dissociated hippocampo-cortical systems vulnerable to each of these two diseases. The implicated genes and proteins may provide promising candidates for more targeted studies of their role in disease-specific neurodegeneration.

### 3.1. Limitations

Our study comes with a number of important limitations that must be addressed. The single greatest limitation of our study is that our gene expression data comes from a limited number of samples taken from only six donors who differed in age, sex and ethnicity. We partially addressed this issue by statistically removing donor effects from our gene expression data and performing leave-one-donor-out analyses, but in doing so, assume certain aspects of gene expression should be fairly consistent across individuals. Some confidence is inspired by the fact that, in spite of these limitations, we were able to replicate findings from rodent studies. We also tried to circumvent this issue by showing that relationships linked to our primary findings hold in several other independent datasets. Another major limitation is a reliance on specific coordinates of samples reported at time of autopsy, translated to single-subject MRI space, and then normalized to a common subject space. While we took measures to improve the quality of the normalization to common space, we cannot rule out noise introduced during any of these steps. Our analysis of gene expression gradients along the hippocampal longitudinal axis is particularly sensitive to these issues because it relies on the exact coordinates of the samples extracted. Once again, we were able to replicate findings from other studies, but it is possible that the importance of some proteins to our model could have been affected. As discussed previously, another limitation is related to our attempts to extrapolate biological importance from machine learning models. While we took many steps to try to test the stability of weights in our models, our interpretations remain somewhat speculative and must be replicated in more focused studies. A recent study suggested that, while the principal community structure of mouse hippocampal connectivity is organized across its longitudinal axis, higher resolution analysis suggests a more complex division of substructures distributed across subfields [8]. We acknowledge that a simple linear gradient may not be sufficient to capture the full complexity of functional organization of the hippocampus, and that this complexity may be driving the variation in our predictions across different hippocampus subfields. Finally, a major limitation comes with the complexity of drawing conclusions across so many datasets, each of which are subject to variation based on methodological processing. We tried to overcome this by primarily using open-access data preprocessed beforehand by experts, and by making all of our data and code freely available at https://github.com/illdopejake/HippocampusAPAxis so that other researchers can scrutinize, reproduce, and hopefully re-use our analyses.

## 4. Acknowledgements

We would like to thank David Berron, Mallar Chakravarty, JB Poline and Gabriel Devenyi for advice and technical support. We would also like to acknowledge support from the Ludmer Centre for Neuroinformatics and Mental Health and the Healthy Brains for Healthy Lives initiative. Additionally, we thank Bruce Miller, Howie Rosen and Bill Jagust for supporting the FDG studies in AD and FTD, which were funded through the following sources: NIA R01 AG045611 (Rabinovici), P50 AG23501 (Miller, Rosen, Rabinovici), P01 AG019724 (Miller, Rosen, Rabinovici). Author J.W.V. is funded through a Tri-Council Vanier CGS Doctoral Fellowship. Author R.L.J. is funded through an Alzheimers Association Research Fellowship (AARF-16-443577, PI: Renaud La Joie). Author R.V.D.W. was funded through a Savoy Foundation fellowship. Nearly all data and tools used in this manuscript are publicly accessible, and this work would not be possible without the hard work put into collecting, storing and maintaining these data and platforms. Data were provided in part by OASIS: Cross-Sectional: Principal Investigators: D. Marcus, R, Buckner, J, Csernansky J. Morris; P50 AG05681, P01 AG03991, P01 AG026276, R01 AG021910, P20 MH071616, U24 RR021382. Data were additionally provided in part by the Brain Genomics Superstruct Project of Harvard University and the Massachusetts General Hospital, (Principal Investigators: Randy Buckner, Joshua Roffman, and Jordan Smoller), with support from the Center for Brain Science Neuroinformatics Research Group, the Athinoula A. Martinos Center for Biomedical Imaging, and the Center for Human Genetic Research. 20 individual investigators at Harvard and MGH generously contributed data to GSP Open Access Data Use Terms Version: 2014-Apr-22 the overall project. Data were additionally provided from numerous studies included in Neurosynth.

## 5. Author Contributions

J.W.V., M.J.G. and R.L.J. conceptualized the study. J.W.V., A.D.P, A.D., E.V.P., C.L., R.V.D.W and B.B. designed and developed the methodologies. J.W.V. analyzed the data. R.L.J. and G.D.R. provided patient data. J.W.V., R.L.J., M.J.G. and A.D.P wrote the manuscript. All authors revised the manuscript and provided critical feedback. R.L.J., M.J.G. and A.C.E. supervised the study.

## 8. Online Methods

All data and analyses described in this manuscript are available online and can be fully reproduced using exclusively open-access software, with (mostly python) scripts and data provided at https://github.com/illdopejake/HippocampusAPAxis. All code and analyses are presented in a series of Jupyter notebooks at the link provided. Supplementary Table S6 outlines which notebook contains the analyses described in each Methods subsections detailed below. See Supplementary Table S5 for a summary of datasets used.

### 8.1. Human gene expression data

Human gene expression data were downloaded from the Allen Human Brain Atlas (http://human.brain-map.org, RRID: SCR 007416). A detailed description of this dataset can be found elsewhere [50, 6]. Briefly, tissue samples were extracted across both hemispheres of two human brain donors, as well as the left hemisphere of four additional donors, totaling 3702 samples. Stereotaxic coordinates and MNI space coordinates are provided for each sample. Each sample underwent microarray analysis and preprocessing to quantify gene expression across 58,692 probes. This analysis provides an estimate of the relative expression of different proteins (encoded by different genes) within the tissue sample. While previous publications have used different strategies to reduce the number of probes (see [6] for review), due to assumptions associated with these strategies and the high-dimensionality approach of our models, we opted to retain all 58,692 probes for analysis.

Importantly, the MNI coordinates originally supplied with the dataset did not account for nonlinear deformations in transforming the donor MRIs in native space to MNI space, and thus included a noticeable degree of error (i.e. many samples mapped outside of the brain or their labeled brain regions) [6]. However, these coordinates have been meticulously reconstructed and trans-formed accounting for nonlinear deformations (http://doi.org/10.5281/zenodo.2483290). Moving forward, all mentions of MNI coordinates will refer to the Devenyi coordinates.

Given the different ages, sexes and other characteristics, substantial differences in gene expression are expected between donors. However, similar to previous studies using this dataset, we were only interested in common patterns of human gene expression for the present analyses, rather than inter-individual differences. As such, all samples across the six donors were aggregated, the effect of donor was removed from each probe using linear models (i.e. with dummy coded donor ID variables), and probe values were standardized. Therefore, probe values represent gene expression normalized across all samples, with inter-individual differences removed.

Along with coordinates, each sample contains ground-truth information about the specific brain sub-structure from which the sample originated, as defined by the anatomist extracting the sample. To identify samples falling within the hippocampus, we selected all samples with structure labels of CA1 field, CA2 field, CA3 field, CA4 field, Subiculum and Dentate Gyrus, from both the left and right hemispheres – 188 samples in total. 18 samples had MNI coordinates more than 3mm outside of the hippocampal volume defined below, leaving 170 hippocampal samples in total.

### 8.2. Identifying the longitudinal axis of the hippocampus

Many previous studies have explored differences between the dorsal and ventral (or posterior and anterior) hippocampus, but such a system requires an often arbitrary delineation between these two structures [20, 42]. To overcome this limitation, we instead sought to quantify the longitudinal axis of the hippocampus and observe changes in gene expression across this axis. Such an approach would still capture gross differences in expression between anterior and posterior sites, but would also allow for detection of more complex gradients. Notably, the hippocampus curves dorsally and medially, so a straight line may not be appropriate for defining its longitudinal structure.

The objective is to identify a curved path that follows the center of mass of the hippocampus along its curvilinear shape **(Fig. S1B)**. The initial hippocampus volume was defined as labels 9 and 19 from the Harvard-Oxford-sub-maxprob-thr25-1mm atlas derived from the MNI ICBM152 average brain template, supplied with FSL 5.0 (RRID:SCR 002823). A “skeleton” of the hippocampal volume was created from morphological operations (dilations/erosions) using the MINC Toolkit (version 1.0.08) (RRID:SCR 014138; http://bic-mni.github.io/#MINC-Tool-Kit). The hippocampus mask was resampled to 0.5mm isotropic voxel size and a chamfer map was created, measuring the distance from the border of the resampled hippocampus volume up to 10mm away. This chamfer map was binarized to create a large smooth blob around the hippocampal surface. An opposite chamfer map was created inside the blob, and the local minimum of the derivatives of this map were computed in order to isolate the points at the greatest distance from the blob surface. This creates a “skeleton” following the curvilinear shape of the hippocampal volume, which was then masked with the original hippocampal volume. Finally, the skeleton was resampled back to 1mm space.

Next, this hippocampal skeleton, in MNI space coordinates, was used to calculate the position of each hippocampus tissue sample along the longitudinal axis. For each sample, we identified the skeleton MNI coordinate with the minimum projected distance to the sample’s MNI coordinate. The position of the sample was then coded as the y-coordinate (anterior-posterior axis) of the closest skeleton voxel. This process effectively transforms all sample coordinates along a single anterior-posterior dimension. **(Fig. S1B)**. Note that, depending on location of the sample, the MNI y-coordinate of the sample may not share the same y-coordinate of the closest skeleton point.

### 8.3. Identifying genes regulating the longitudinal axis of the human hippocampus

We sought to identify which genes may play a significant role in the positioning of samples along the longitudinal hippocampal axis. Sparse regression algorithms built for high dimensional datasets have been proposed, such as least-angle regression (LARS) and LASSO-LARS. However, during regularization, these algorithms will often select only one of a set of several collinear variables and reduce the coefficient of the other variables in the set to zero. In the case of gene expression data, gene co-expression networks are of interest to us, and we do not necessarily want to select one of a set of co-expressed genes. Therefore, we opted instead to use a LASSO-PCR approach [53, 30]. Such an approach will reduce the dimensions of the data while preserving gene co-expression networks, yet still allow for a sparse selection of features.

In summary, we reduced our input data, a 170 (sample) x 58,692 (probe) matrix, using principle components analysis (PCA) with singular value decomposition. The resulting 170 (sample) x 170 (component score) matrix was used in a principal component regression (PCR) model **(Fig. S1)**. Approaches to PCR models typically reduce the number of independent variables by removing the components whose eigenvalues fall below some threshold related to the percentage of variance explained. This does not account for potentially strong relationships between the dependent variable and minor components. Thus, we elected to use a Least Absolute Shrinkage and Selection Operator (LASSO) regression model with sample position along the longitudinal hippocampus axis (defined in the previous section) as the dependent variable.

In our regression model we have our standardized matrix of gene expression data *X*, our measurements along the longitudinal axis *Y*, and the model *Y* = *XB* + *ϵ*. We wish to estimate the values of the matrix *B* = [*β*_0_*, β*_1_*, …, β_p_*]*^T^*, where *β_i_* is the estimated impact of probe *i* on longitudinal position. Probes with larger impacts will have higher estimated values; negative values suggest greater expression in posterior compared to anterior hippocampus, and vice versa. Since there are a large number of regression parameters, we use dimension reduction through PCA. We transform the data such that *X^T^X* = *P* Λ*P^T^* = *Z^T^Z*, where Λ is the diagonal matrix of eigenvalues of *X’X*, *Z* is the matrix of principal components, and *P^T^P* = *I*. We are now interested in solving the principal component regression *Y* = *ZA*, where the regression coefficients are stored in the matrix *A* and are the contribution of principal components to position. We derive estimates of *A* using LASSO. The coefficients of the two regression equations are related by the expressions *A* = *P^T^B* and *B* = *P A*, so we estimate 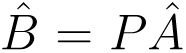, giving us the beta values of the individual probes, which are in terms of the original probes.

There are limitations to this approach. Beginning with the full set of components can incidentally retain small components and make estimates of beta coefficients unstable [30]. Interpretation of the components is challenging, and here they were generated without the dependent variable (the measurements along the anterior-posterior axes). At the theoretical level PCA can break down when there are many more variables than observations since the sample covariance eigenvectors may not be close to population eigenvectors [24] though empirical results here are positive and in concordance with previous results. Partial least squares (PLS) is a method related to PCR that accounts for the dependent variable and returned similar results (Figure S4).

To test the generalizability of the model, we employed several cross-validation methods. First, we performed 10-fold cross-validation of the full data set, which was repeated 10 times. Second, we performed a leave-one-subfield-out cross-validation, to see if a model defined on five hippocampal subfields (CA1-4, subiculum, dentate gyrus) could predict the axis position of samples from the sixth subfield. Finally, we performed leave-one-donor-out cross-validation to see if a model trained on samples from five donors could predict the axis positions of samples from the sixth donor. Note that the range of sample position was constrained by anatomy during the leave-one-subfield-out cross-validation, and the number of samples varied quite dramatically across donors for the leave-one-donor-out validation. The final model used for all subsequent analyses utilized all samples. As a sanity check, we calculated the mean of the fifty genes with the highest (anterior) and lowest (posterior) betas within each sample and measured the variance explained in sample position along the longitudinal axis by this average expression signal.

### 8.4. De-constructing model features to assess candidate genes responsible for axis regulation

An advantage of the LASSO-PCR model is that it is more likely to identify several genes participating in a co-expression network rather than arbitrarily identifying a single gene to represent that network. However, this also leads to a possible disadvantage related to reduced precision in singling out which genes, if any, are singularly important to the model. Additionally, the global feature importances of a LASSO model cannot be reliably interpreted, as adding or removing features can cause feature importances to shuffle dramatically [25]. We attempted to de-construct our model with these limitations in mind. Fifty probes with, respectively, highest (anterior) and lowest (posterior) back-transformed weight (feature importance) were iteratively removed from our model. After each removal of these 100 probes, the model was refit, 10-fold cross-validation (CV) accuracy was recorded, and the 100 top probes from the new model were removed. This process was repeated until all probes were removed. As a control, we repeated this same process iteratively removing 100 random probes instead of the 100 most important probes. Change in CV accuracy across rounds of probe removal was visually assessed and inflection points were identified at rounds where CV accuracy dropped and did not recover. Rounds in between inflection points were considered stable, and probes removed between inflection points were grouped together in gene sets, analyzed separately in subsequent analysis.

To establish whether these gene sets alone could predict sample position along the longitudinal axis of the hippocampus, we reran the LASSO-PCR model with only the probes involved in these gene sets. Prediction accuracy was recorded using 10-fold cross-validation. The models were run ten times with bootstrap samples to attain confidence intervals. As a control analysis, models were run using sets of random probes the same size as each gene set, and this process was repeated 10 times for each set, each time using cross-validation to measure prediction accuracy. Finally, in order to compare larger gene sets to Set 1 – which contained only 100 probes – we extracted 10 random sets of 100 genes from within each gene set and input these into the model, once again using 10-fold cross-validation to measure prediction accuracy.

To further highlight candidate genes involved in hippocampal longitudinal axis regulation, we employed the Local Interpretable Model-Agnostic Explanations (LIME) python package (https://github.com/marcotcr/lime/). LIME makes local perturbations to model inputs and measures the impact of those perturbations on model performance. LIME can only assess local feature importance, however, by aggregating information across multiple local features, some limited information can be ascertained about contribution of features (probes) to predicting an outcome (sample position along the longitudinal axis). For each gene set identified, we performed 10-fold cross-validation with a Random Forest Regressor. A Random Forest Regressor was chosen because its metric of feature importances is itself assessed using out-of-sample prediction. For each fold, LIME was used to identify absolute feature importances for samples in the left-out fold, and this information was aggregated across all predictions from all folds. Elevated feature importance could indicate importance of a probe across prediction of multiple samples, or could indicate great importance across a limited set of predictions, meaning interpretation is still limited.

### 8.5. Characterization of gene sets using gene ontology enrichment analysis

Gene ontology (GO) enrichment analysis was used to characterize functions shared by several genes within gene sets. These analyses were performed using the online tool GOrilla (RRID:SCR 006848; http://cbl-gorilla.cs.technion.ac.il/), which identifies terms from the GO libraries that are associated with genes in the inputted gene set and are significantly (FDR < 0.1) enriched compared to a baseline gene set. We used the entire set of genes available in the Allen Human Brain Atlas dataset as the baseline gene set. Altogether, the background set we entered included 29,381 distinct genes, 19,895 of which were recognized by GOrilla. Of these, only 17,836 were associated with a GO term. We left all other parameters to their defaults. Some of the gene sets produced long lists of enriched terms. We summarized this information using hierarchical agglomerative clustering on the significantly enriched terms. A binary gene x term matrix was created where a 1 indicated a gene was associated with a term. This matrix was fed to an Agglomerative clustering algorithm using Jaccard index with average linkage and pre-calculated connectivity constraints (10 neighbors), and the process was repeated varying the number of clusters from 2-20. Local peaks in silhouette index were used to define the final cluster number, favoring a higher number of clusters for better precision. The resulting clusters represented sets of genes sharing several associated terms. For gene Set 2 (top 101-600 most important probes to the model, see Section 2.2), peaks in Silhouette score were seen at k=2 (0.225), k=7 (0.132) and k=10 (0.128). We chose a 10-cluster solution. For gene Set 3 (top 601-2700 probes, Section 2.2), peaks in Silhouette score were seen at k=2 (0.349), k=5 (0.173) and k=12 (0.093). We chose a 12-cluster solution. The purpose of this analysis was to cluster genes with enriched GO terms for purely descriptive purposes.

### 8.6. Whole-brain genomic representation of the hippocampal longitudinal axis – HAGGIS formulation

We sought to ascertain to what degree the specific pattern of genes regulating the hippocampal longitudinal axis was expressed throughout the rest of the brain. The probe weight (beta) vector from the LASSO-PCR analysis can be thought of as a hippocampal longitudinal axis genomic signature. In order to test for the presence of this signature in other brain regions, we found the dot product between the beta vector (genomic axis signature) and the gene expression (probe) vector for each sample **(Fig. S1C)**. Note that when estimating regression coefficients we have:

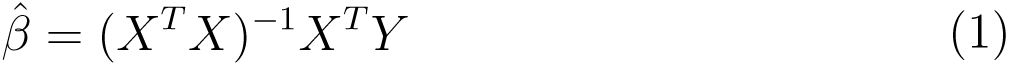

This is equivalent to using the estimates of coefficients from the LASSO-PCR model to predict the location of the (non-hippocampal) sample along the hippocampal axis. In practice, this amounts to using the hippocampus model to predict where a non-hippocampus sample might fall along the hippocampal longitudinal axis based on that sample’s gene expression. However, conceptually, this value can also offer an index of covariance between a given sample’s gene expression and the gene expression profile of the anterior or posterior hippocampus. Higher (positive) values represent greater genomic covariance with the anterior hippocampus, while lower (negative) values represent greater similarity to the posterior hippocampus. For the purposes of parity, this index will be referred to in the text as the Hippocampal Axis Genomic Gradient Index of Similarity (HAGGIS) index.

### 8.7. Comparisons with resting-state functional connectivity

For each of the 170 hippocampal samples, a resting-state functional connectivity map was downloaded from Neurosynth (RRID:SCR 006798; http://neurosynth.org/) using the closest available MNI coordinate to the MNI coordinate of the sample. The Euclidian distance between Neurosynth coordinate and sample coordinate never exceeded 2mm. Each map is based on the resting-state functional connectivity patterns of 1000 young, healthy individuals from the Brain Genomics Superstruct project [55].

We sought to test whether the genes regulating the longitudinal axis of the hippocampus contribute to the differential brain connectivity observable along this axis. The measurement resolution of resting-state functional magnetic resonance imaging (rsfMRI) limits detail at which differences in connectivity can be observed along a structure as small as the hippocampus. To ameliorate this issue, we divided the hippocampus into genomically-determined posterior and anterior subsections, created mean connectivity maps for each, and used these mean connectivity maps to create a subtraction image representing differential functional connectivity between the two poles of the hippocampus [28]. To determine a reasonable division between anterior and posterior hippocampus, we created a split at every point along the hippocampus skeleton. For each split, we classified samples as anterior or posterior based on the position of the coordinate along the longitudinal axis relative to the split. For each split, we next ran Logistic Regression, entering sample class (i.e. anterior or posterior) as the dependent variable and sample HAGGIS as the only independent variable. We then plotted the classification accuracy at each split under the hypothesis that higher anterior-posterior classification accuracy would suggest a more empirically sound anterior-posterior division (Fig S4A). We defined the optimal anterior and posterior cut points as i) local maxima in accuracy that ii) were at least 3mm from both hippocampal poles and iii) captured at least 20 samples for each side of the split. This lead to an anterior split point of y=108 (MNI: −19) and a posterior point of y = 94 (MNI: −35). All samples in between were removed. Results in the main text are reported using this split but, due to the somewhat arbitrary nature of this analysis, results are also reported for several other splits.

Once the anterior and posterior samples had been defined, a mean image was made of the functional connectivity maps corresponding to each anterior and posterior sample, respectively. The posterior map was then subtracted from the anterior map. The resulting image represented relative functional connectivity to the anterior hippocampus over the posterior hippocampus. For each non-brainstem, non-cerebellum sample, a 5×5×5mm cube was drawn around the MNI coordinate of the sample. The mean of rsfmri subtraction image values within the cube was calculated, and this value was used as a measure of relative functional connectivity of the sample to the anterior over posterior hippocampus. Finally, we ran a Pearson’s correlation between this functional connectivity measure and the HAGGIS. A positive correlation would indicate that brain regions with more genomic similarity to the anterior or posterior hippocampus would be more likely to be functionally connected to those regions, respectively. This analysis was performed using weights from the model performed on the entire gene set, as well as weights from models defined on individual gene sets.

We repeated this analysis using three other brain masks: i) All brain regions; ii) All regions except cerebellum, brainstem and hippocampus; iii) cerebral cortex only. In addition, we varied the radius of the cube drawn around the sample coordinate between 1mm and 6mm. For completeness, we performed the above analysis using each cube radius, with each mask, and using many different splits – a total of 336 analyses. To ensure the relationships between HAGGIS and rsfMRI connectivity were not born out of chance, we performed a permutation test for each of the 336 conditions. Specifically, the gene expression values for each sample were randomly shuffled, and a correlation was run between the extracted rsfMRI connectivity values and the shuffled gene expression values. This process was repeated 1000 times to create a null distribution, to which the observed value was compared to establish an exact p-value.

We performed one final validation by applying diffusion map embedding [36, 52] – a non-linear dimension reduction approach – to the hippocampal-brain functional connectivity matrix. This approach summarizes variation in hippocamus-brain connectivity into components or “gradients” [52], allowing threshold-free representations of variation in hippocampus-brain functional connectivity for each tissue sample. The whole-brain connectivity maps for each sample (see above) were masked with a cortex-only mask (see above), vectorized and concatenated into a Sample x Voxel matrix. A correlation matrix was created from the transpose, generating a Sample x Sample similarity matrix, which was reduced using diffusion map embedding with default settings. We report the total variance in hippocampus-brain functional connectivity explained by each gradient, as well as the r^2^ summarizing each gradient’s relationship to sample location along the longitudinal axis, and predicted sample location based on gene expression (proportionate to HAGGIS). We also report p-values, which are Bonferroni corrected for multiple comparisons. We then selected the gradient with the greatest relationship to predicted sample location (i.e. HAGGIS), provided this relationship was significantly stronger than that of other gradients, as measured using Steiger’s tests [52]. For these select gradients, we also report this information with sample location predicted using each of the gene Sets described above (Section 2.2, 8.4).

Other studies have published examining genomic regulators of functional connectivity [47, 54], and so we sought to understand what proportion of the variance explained from the main analysis (shown in Fig 4A) was unique to the HAGGIS rather than general network connectivity. We trained a cross-validated PLS model to learn the genomic features predicting relative anterior vs posterior connectivity to the hippocampus (i.e. the map in Fig 4A; see subsection 8.10 below for details). We considered the 10-fold cross-validated variance explained of this model to represent an estimate of the maximum variance explainable given the present genomic data. We then represented the variance explained of HAGGIS as a proportion of the overall variance explainable given the genomic data (visualized in Fig 4C).

### 8.8. Comparisons with structural covariance

Structural covariance is thought to reflect shared cytoarchitecture and/or developmental and degenerative trajectories between regions [2]. The anterior and posterior hippocampus have shown different patterns of structural covariance with the rest of the brain [39], and structural covariance appears to be genetically determined to some extent [2]. Accordingly, we assessed whether the differential structural covariance between different brain regions and the hippocampus along its longitudinal axis is reflected by patterns of genomic covariance.

Structural covariance was calculated using the OASIS: Cross-Sectional structural (T1) MRI dataset [35], accessed with Nilearn (RRID:SCR 001362; https://nilearn.github.io/). The OASIS images came preprocessed using the SPM DARTEL pipeline [7]. 153 preprocessed gray matter volume images were identified as healthy, cognitively normal young (age < 40) controls. For each voxel corresponding to the MNI coordinates of an Allen Human Brain Atlas hippocampus sample, a structural covariance vector was calculated between that voxel and all other brain voxels. Elements in the vector represented Pearson correlation coefficients between voxel values across the dataset of 153 individuals between the two regions. Anterior and posterior hippocampus divisions identified in the previous analysis were used to divide the covariance vectors, and the average covariance within anterior vectors and posterior vectors were calculated, respectively. The difference between these vectors was calculated to create a map where each voxel contained a value representing the relative structural covariance to the anterior over the posterior hippocampus. The values strongly favored the anterior hippocampus, so the map was z-scored, such that lower values represented less structural covariance to the anterior hippocampus. Relationships between HAGGIS and relative structural covariance were carried out in a manner identical to the functional connectivity analysis described above, and were repeated using different gene sets and brain masks. Similar to the functional connectivity analysis, we calculated the variance explained by HAGGIS as a proportion of the maximum variance explainable given the data (see previous subsection).

As with the functional connectivity analysis, we used diffusion map embedding to generate threshold-free measures (gradients) summarizing hippocampus-brain structural covariance. For each sample, we calculated structural covariance between the voxel at the sample location and all other voxels falling within in a cortical mask, creating covariance vectors. These vectors were concatenated into a Sample x Voxel matrix, and reduced using diffusion map embedding as described above (Section 8.7).

### 8.9. Comparisons with neurodegeneration in Alzheimer’s disease and frontotemporal dementia

Previous studies have noted the differential relationship of the hippocampus to Alzheimer’s disease (AD) and frontotemporal dementia (FTD). We tested whether regions more genomically similar to the anterior than posterior hippocampus might be more vulnerable to neurodegeneration in FTD than in AD (and vice versa). In April 2018, we queried our database looking for patients who fulfilled the following criteria: i) Had available both a [^11^C] Pittsburgh Compound B (PiB)-PET scan for *β*-amyloid and a [^18^F] Fluorodeoxyglucose (FDG)-PET scan of brain glucose metabolism acquired on the Biograph scanner; ii) Had either a clinical diagnosis of AD [37] and a positive PIB-PET read, or a clinical diagnosis of FTD (either behavioral variant FTD or semantic variant primary progressive aphasia, as described in [38]) and a negative PIB-PET read. Note that This query resulted in 36 AD and 39 FTD patients. Five patients were later excluded because of incomplete FDG-PET SUVR (missing at least one of the 6 frames between 30 and 60 min post-injection), resulting in a final count of 35 AD and 35 FTD patients. Demographic information can be found in Table 2. Note there is no overlap between this sample and the sample described in [28].

**Table 2:**
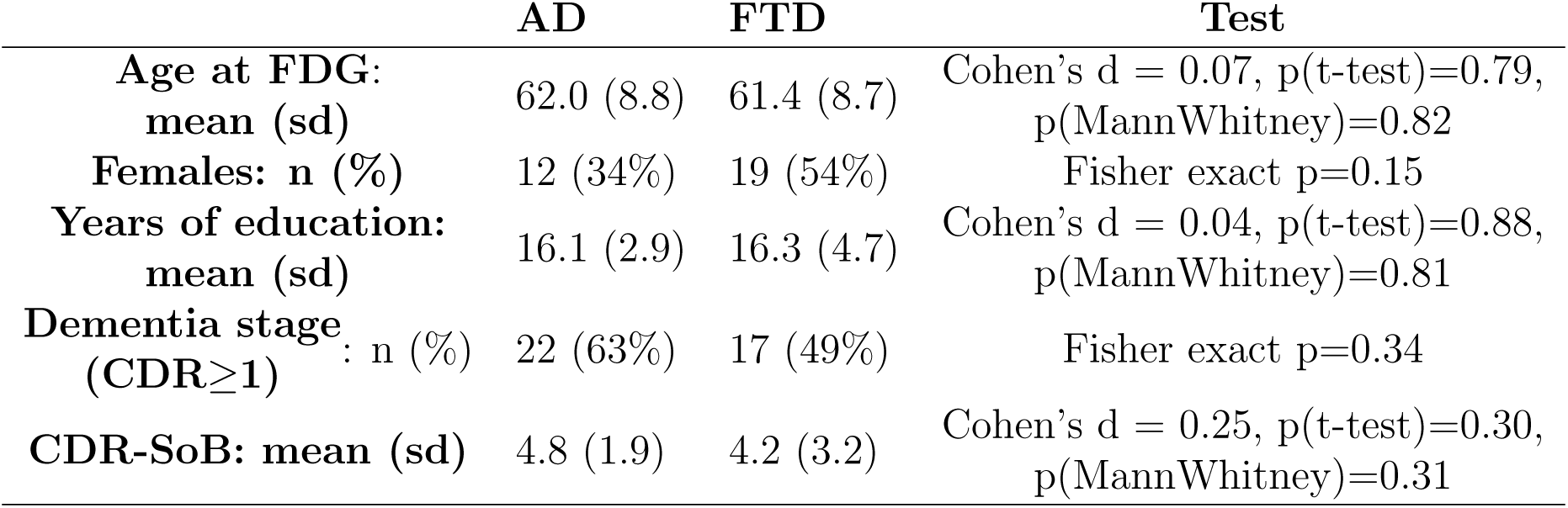
FDG = fluorodeoxyglucose; sd = standard deviation; CDR = Clinical Dementia Rating; CDR-SoB = Clinical Dementia Rating, Sum of Boxes

All patients were seen at the at University of California, San Francisco Memory Aging Center and imaged at the Lawrence Berkeley National Labs. PET acquisition details can be found elsewhere [40]. FDG-PET images were processed using SPM12 using a previously described pipeline [40]. Briefly, six five-minute frames were realigned and averaged, and the average image was coregistered onto patient specific anatomical T1-MRI scans. Standard uptake value ratios (SUVR) were calculated using the pons (Freesurfer segmentation of the brainstem with manual cleaning) as a reference region, and SUVR images were warped to the MNI template using MRI-derived parameters. All 70 patients were entered into a voxelwise t-test controlling for age and disease severity (Clinical Dementia Rating Sum of Boxes score) using SPM12, highlighting differences in glucose hypometabolism (a proxy for neurodegeneration) between AD and FTD patients. The t-map from this analysis was used for subsequent analyses, and is made available with this publication (https://neurovault.org/collections/4756/).

For each non-brainstem, non-cerebellar sample, a 5mm diameter cube was drawn around the sample’s MNI coordinates, and the mean t-value from the t-map described above was extracted. This value represents the relative neurodegeneration in FTD over AD in or around the region the sample was extracted from. Across samples, a correlation was calculated between this value and the sample’s HAGGIS. A positive correlation would suggest regions more genomically similar to the anterior than the posterior hippocampus are more vulnerable to neurodegeneration in FTD than in AD. To ensure our findings were not specific to the brainmask used or the size of the extraction cube, we reran the analysis using each of the three additional masks described in Section 8.7, as well as varying the diameter of the extraction cube. Finally, permutation tests were run for each condition to compare our observations to chance (see Section 8.7). As with the previous analyses, we ran these sets of analyses across different gene sets.

### 8.10. Identifying candidate genomic regulators of brain-hippocampus interactions

In sections 8.7, 8.8 and 8.9, we describe methods to uncover relationships between HAGGIS and hippocampus-brain interactions. We wished to identify which specific genes were principally involved both in the organization of the longitudinal axis of the hippocampus, as well as in the hippocampus-brain interactions, further elucidating the role of the various genes identified in section 8.4 along the axis. For each hippocampus-brain interaction map (visualized in Fig. 4A), we fit a partial least squares (PLS) regression model with gene expression information as X and hippocampus-brain interaction value as Y, across all brain samples. As with the model described in section 8.3, the X input was first transformed using principal components analysis and represented as a set of genomic components. The model was fit varying the number of PLS components (i.e. modes) between 1 and 10, and using 10-fold cross-validation to assess model accuracy. The model with the highest cross-validated explained variance was selected as the best model, and was considered the maximum explainable variance given the genomic data available, which was therefore useful to compare to the HAGGIS models (see section 8.7 above). Note that the hippocampus itself was not included in any of the models. For each of the three PLS models, feature weights were backtransformed back into probe space (see section 8.3), and the top 50 anterior and posterior associated features (i.e. with the highest and lowest weights) were identified. Overlapping features between each model and the hippocampus longitudinal axis model are reported. These features represent genes that appear to be very important in predicting the location of tissue samples in the hippocampus, but also in predicting interactions between the hippocampus and other brain regions. To ensure this overlap did not occur by chance, 1000 sets of 100 random probes were generated, and used to calculate the probability of overlap between 100 random features and the 100 features from the the hippocampus longitudinal axis model.

### 8.11. Comparisons with large-scale cognitive systems

The Neurosynth website contains 3D meta-analytic functional co-activation maps from task-fMRI studies that are paired with sets of related topics (words) extracted from the text of these studies. These topic-list/co-activation map pairs are the result of a Latent-Dirichlet Allocation across 11,406 articles, the details of which can be found elsewhere [41]. In short, topic lists represent words that are mentioned greater than chance (FDR<0.01) in papers reporting functional co-activation in given coordinates, summarized by paired co-activation maps. All 100 (association/reverse inference) maps from the set of 100 topic list/co-activation map pairs on the Neurosynth website were downloaded and binarized such that all values above 0 were set to 1, and all other values were set to 0. We manually labeled the topics according to their hypothesized association with the AT-PM system [44] based on the content of the word list (AT/PM/Not associated) but without reference to the spatial pattern of the co-activation. For each of the 100 binarized functional meta-analytic co-activation maps, all samples with MNI coordinates falling within the map were identified, and the mean HAGGIS of those samples was calculated. Therefore, each topic/map pair had an associated value indicating the degree to which the brain regions involved expressed genes similar to the anterior or posterior hippocampus. Higher values represented similarity to the anterior hippocampus, lower values to the posterior hippocampus, and higher absolute values represented greater genomic covariance. To increase confidence in this approach, the main analyses were restricted only to maps overlapping with at least 500 samples (29/100).

To help visualize these results, we created a word cloud summarizing both the spatial (functional coactiviation) and topic (cognitive) information associated with the anterior and posterior hippocampus respectively. For the topic information, each topic-set contained 40 words arranged by importance to the topic-set. Each word was given a value proportionate to its importance rank in its topic set (i.e. most important word valued at 40, least important at 1). Next, the value of each word was multiplied by the average HAGGIS within the binarized map paired to the word’s topic-set (i.e. the bars in Fig 5), multiplied by 1000 to increase the weighting of this multiplier proportionate to the within-set ranking. Therefore, each word had an associated value, such that the highest values represented words most important to topic/map pairs with the greatest HAGGIS, where multiple mentions increased the value of the word. To summarize the spatial information, we binarized each map and multiplied it by the average HAGGIS within the binarized map (i.e. the bars in Fig 5), and summed all maps, and smoothed the image with a 4mm isotropic kernel. All voxels with positive values were binarized into a mask, and this mask was used as constraint for the anterior-hippocampus word cloud, inside which the top 100 words were visualized. All voxels with negative values were binarized into a posterior mask used as a constraint for the posterior-hippocampus word cloud. The word values were repeated inverting the HAGGIS multipliers, and the top 100 words were visualized. The final image represents brain regions coactivated more with the anterior vs posterior hippocampus, and the cognitive topics most associated with those regions.

## 9. Supplementary Figures

**Supplementary Fig. S1:**
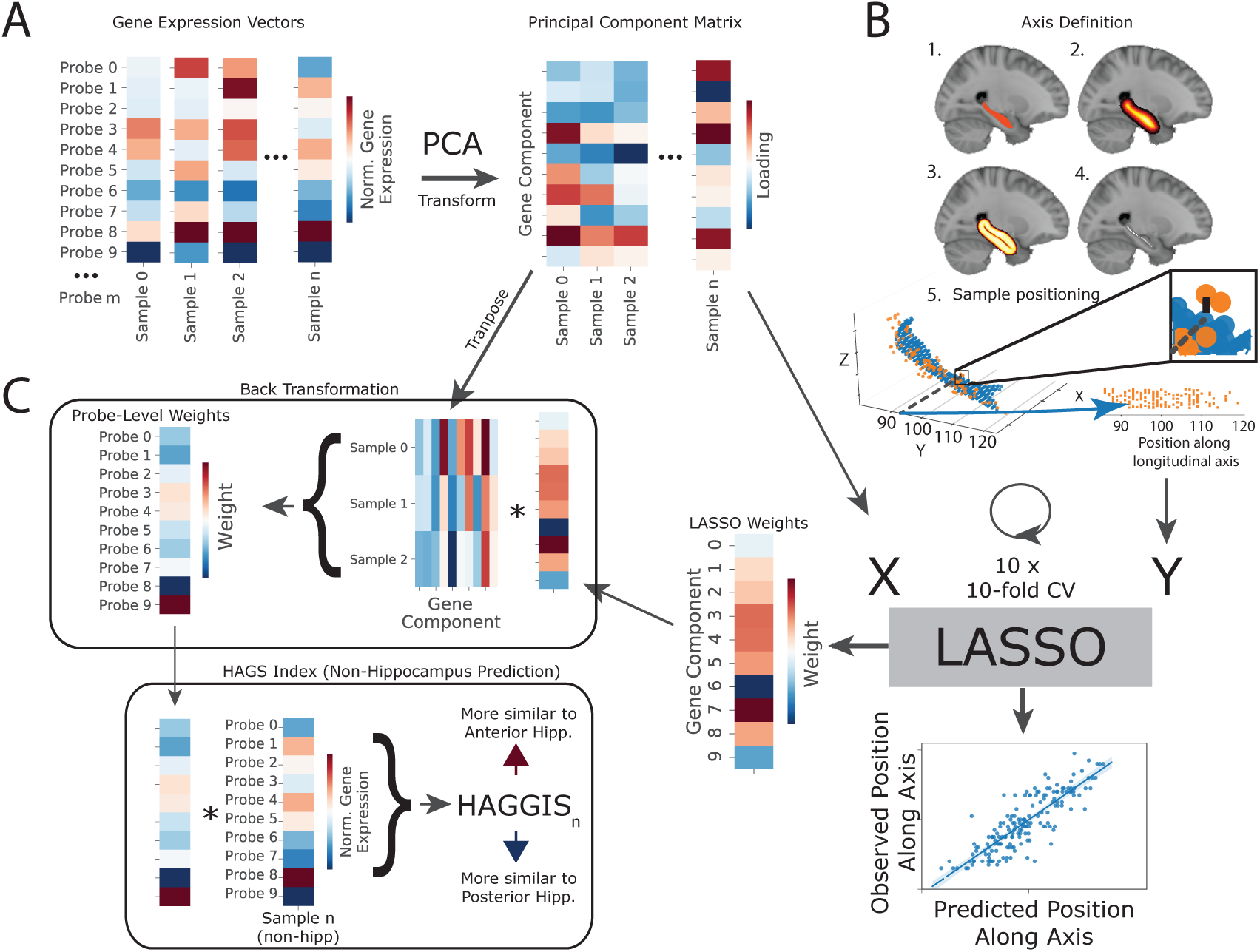
LASSO-PCR pipeline to predict the position of a tissue sample along the longitudinal axis of the hippocampus using gene expression. **(A)** The 170 (Sample) x 58,692 (probe) gene expression matrix was first reduced using principal components analysis (PCA), such that each sample had a singular value representing the loading onto each principal component. The principal component matrix was used as the predictor (X) variable in the LASSO-PCR model. **(B)** The longitudinal axis of the hippocampus was defined with a medial axis transform: 1) We start with a mask of the hippocampus, which is resampled to 0.5mm space. 2) The mask is dilated by creating a chamfer map measuring distance from the center of the hippocampus, extending out 10mm into a smooth hippocampus-shaped blob. 3) An inverse chamfer map was created inside the blob, local minimum of the derivatives of this map were computed. 4) These operations resulted in a hippocampus “skeleton”. 5) For each tissue sample (orange), the closest hippocampus skeleton voxel (blue) was located, and the y-axis of this coordinate was used as the position of the sample along the longitudinal axis, which was used as the dependent variable (Y). **(C)** A sparse LASSO regression model fit the (reduced) gene expression data to position along the atlas, with ten rounds of 10-fold cross-validation. Model weights were back-transformed to probe space. The back-transformed weights were applied to the gene expression vectors of non-hippocampus samples to the derive the HAGGIS, indicating genomic similarity to the anterior (positive) or posterior (negative) hippocampus.

**Supplementary Fig. S2:**
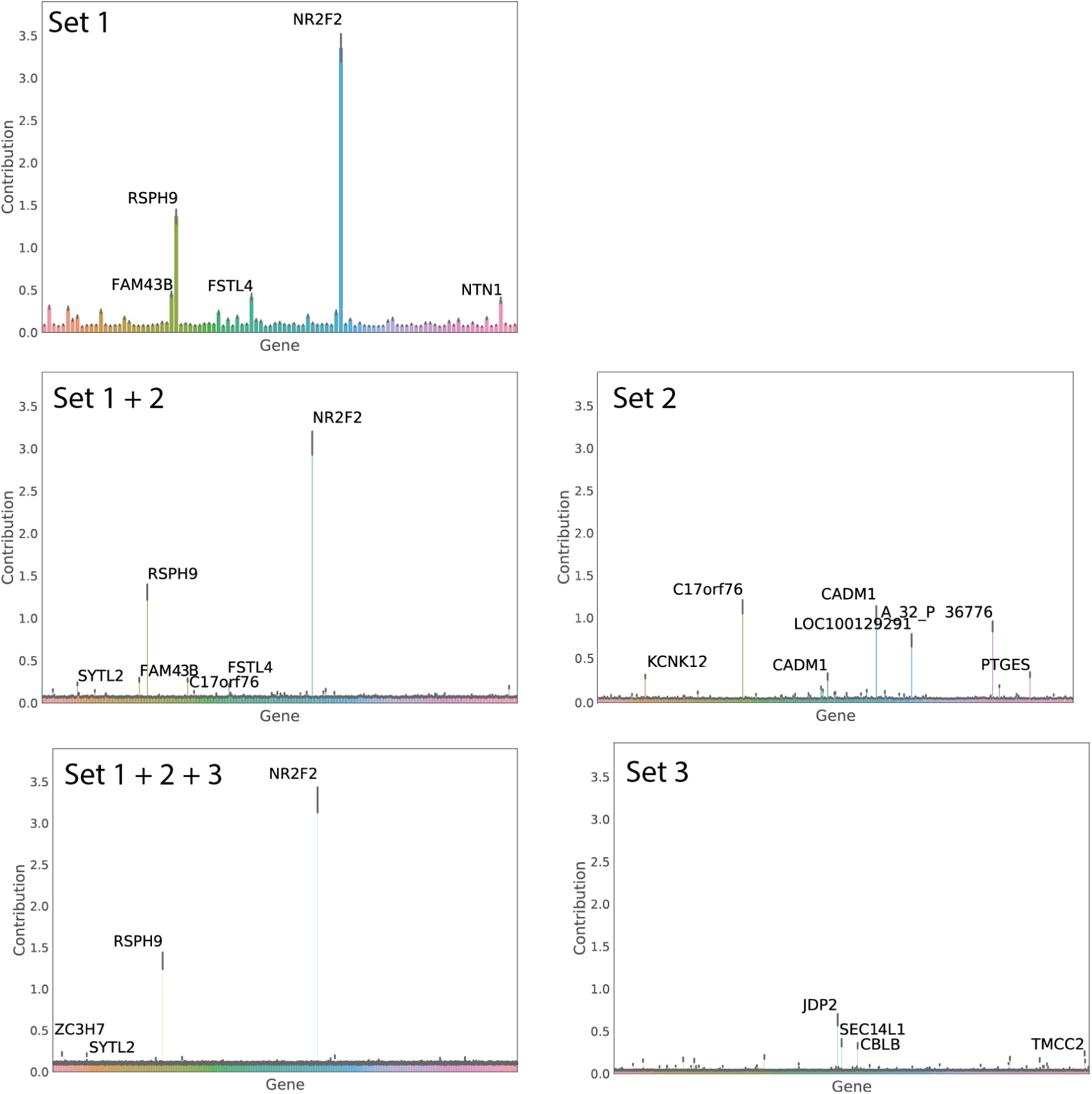
Feature-explainer applied to different gene sets. The Random-Forest based feature explainer was applied to different combinations of gene sets associated with position along the longitudinal axis of the hippocampus. For each plot, the y-axis represents local feature importance, indicating the degree to which, on average, perturbing the feature (probe) impacts individual model predictions. NR2F2 and RSPH9 consistently demonstrated the greatest importance when included in the model. Compared to Set 1, feature explainers identified more features with less importance for Sets 2 and 3.

**Supplementary Fig. S3:**
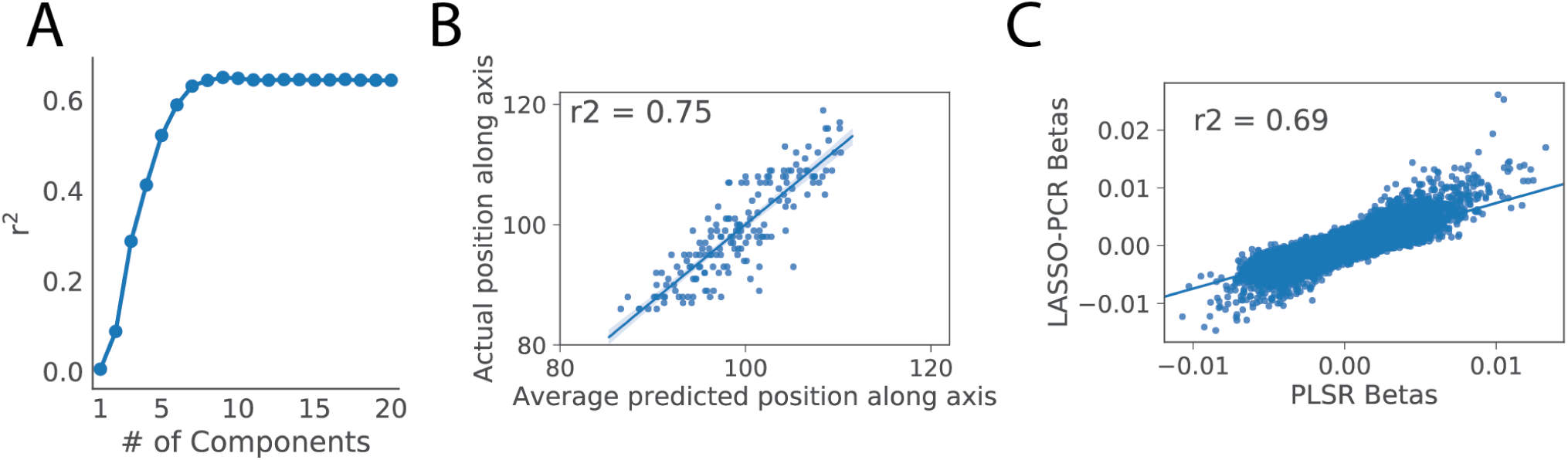
Validating results with PLSR. To ensure previous findings were not a product of algorithm choice, PLSR was fit to the gene expression data in order to predict position along the longitudinal axis of the hippocampus. **A** 10-fold cross-validation suggested nine as the optimal number of components. **B** Fitting the PLSR model to the data resulted in a similar r^2^ as the LASSO-PCR approach. **C** The weights from the LASSO-PCR and PLSR models were highly correlated.

**Supplementary Fig. S4:**
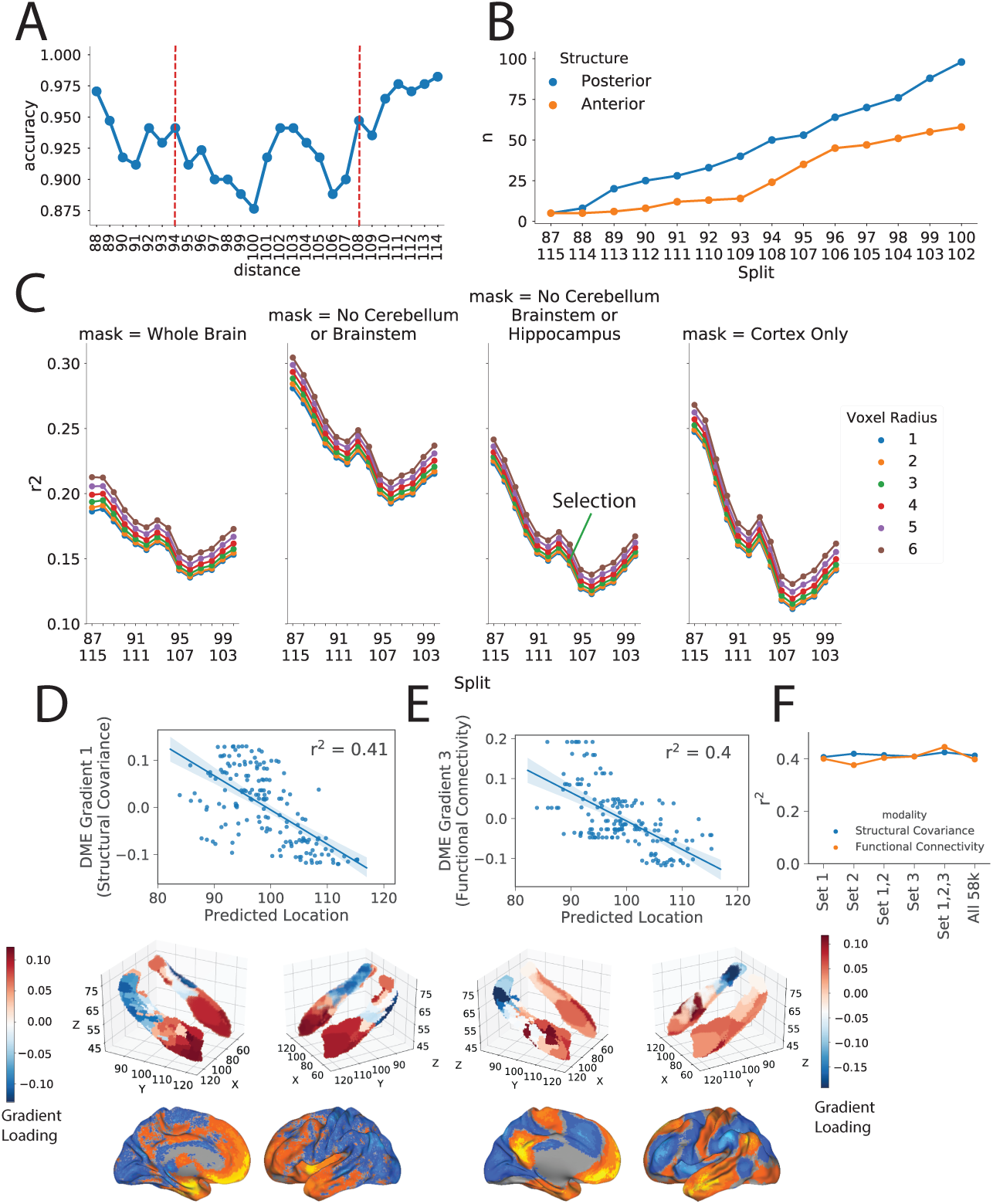
Validation of rsfMRI connectivity results. **(A)** An anterior-posterior split of the hippocampus was made at every y-coordinate along the hippocampal axis, and a Logistic Regression with HAGGIS was performed to classify anterior from posterior hippocampus. Accuracy at each split is visualized. The coordinates of the final split used for the analysis in the main text are indicated with red dashed lines. **(B)** The analysis was performed across several additional splits, indicated on the x-axis. The number of anterior and posterior samples included after each split are shown in orange and blue, respectively. The splits move from more extreme to more central as the x-axis moves from left to right. **C** The rsfmri analysis was repeated varying the radius of the extraction cube, the brain mask, and the anterior/posterior split. The r^2^ of the correlation between HAGGIS and functional connectivity for each condition is shown. Diffusion map embedding was used to summarize principal axes of whole-brain functional connectivity (**D**) and structural covariance (**E**). Select gradients are correlated with the gene expression pattern predicting longitudinal axis location. The gradients are rendered onto a hippocampus surface, and expression of the gradient in whole-brain connectivity/covariance patterns is visualized. **F** The r^2^ of relationships shown in C and D where the gene expression pattern is composed of different gene sets.

**Supplementary Fig. S5:**
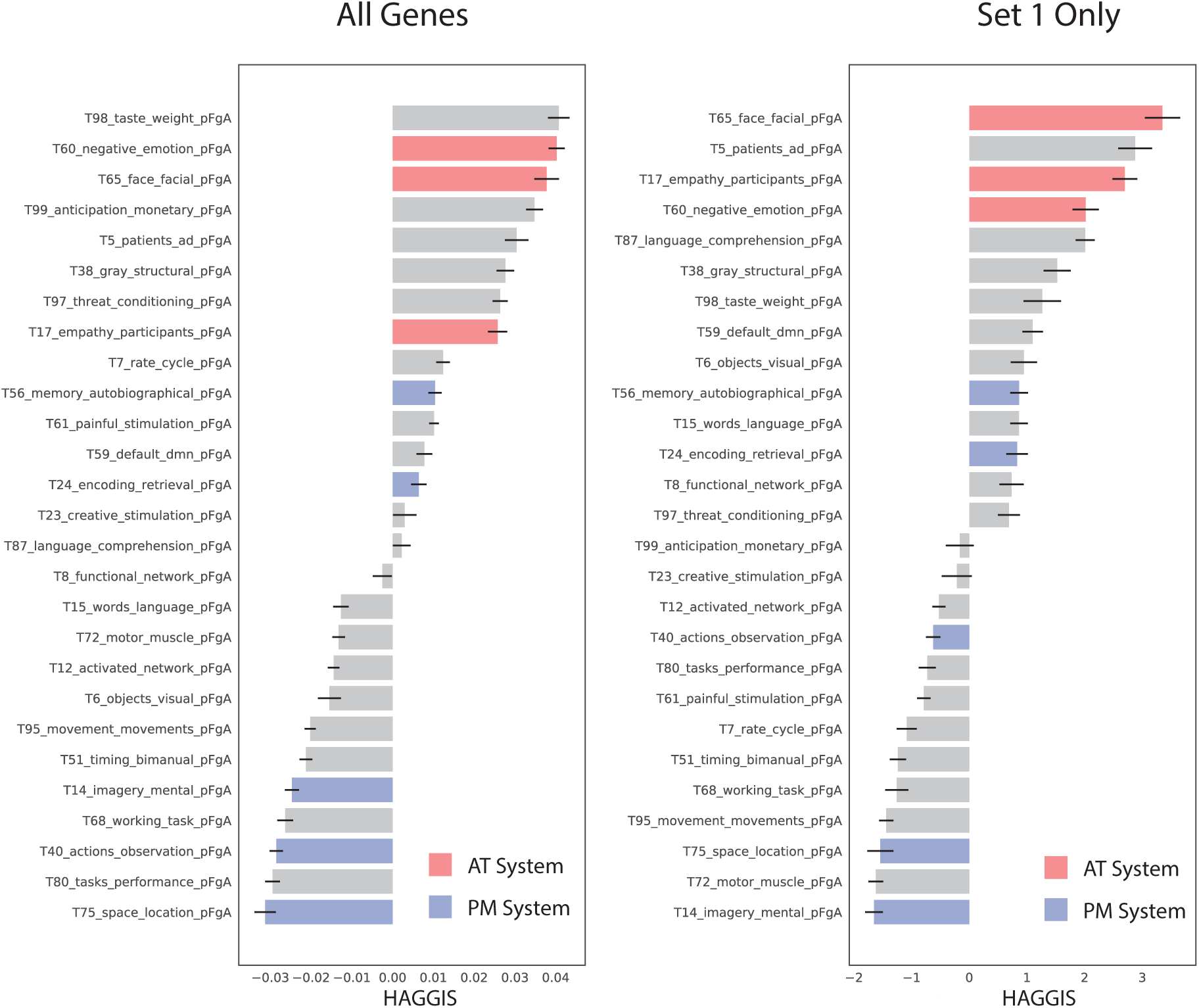
Cognitive meta-analysis when using all probes vs. top 100 probes. On the left is a vertical reproduction of Fig 4F. On the right is the results of the exact same analysis, except calculating the HAGGIS using only the top 100 probes, rather than all 58,692 probes. The pattern is remarkably similar, especially as pertaining to the topics associated with the AT/PM system.

**Supplementary Fig. S6:**
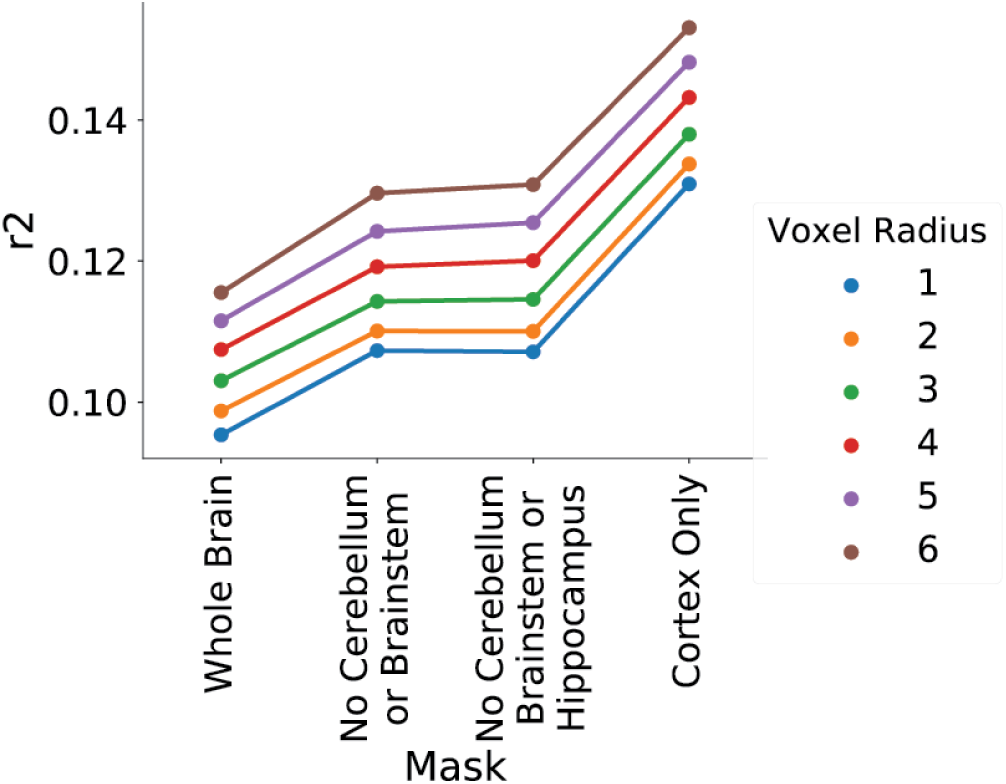
Validation of FDG neurodegeneration results. The analysis comparing HAGGIS to relative neurodegeneration in AD vs FTD was repeated using different extraction cube sizes and different brain masks. The r^2^ for each condition is visualized

## References

[1] Adnan, A., Barnett, A., Moayedi, M., McCormick, C., Cohn, M., & McAndrews, M. P. (2016). Distinct hippocampal functional networks revealed by tractography-based parcellation. Brain Structure and Function, 221, 2999–3012.

[2] Alexander-Bloch, A., Giedd, J. N., & Bullmore, E. (2013). Imaging structural co-variance between human brain regions. Nature Reviews Neuroscience, 14, 322–336.

[3] Alexander-Bloch, A. F., Mathias, S. R., Fox, P. T., Olvera, R. L., Göring, H. H. H., Duggirala, R., Curran, J. E., Blangero, J., & Glahn, D. C. (2017). Human Cortical Thickness Organized into Genetically-determined Communities across Spatial Resolutions. Cerebral Cortex, pp. 1–13.

[4] Alzu’Bi, A., Lindsay, S. J., Harkin, L. F., McIntyre, J., Lisgo, S. N., & Clowry, G. J. (2017). The Transcription Factors COUP-TFI and COUP-TFII have Distinct Roles in Arealisation and GABAergic Interneuron Specification in the Early Human Fetal Telencephalon. Cerebral Cortex, 27, 4971–4987.

[5] Andersen, P., Morris, R., Amaral, D., Bliss, T., & O’Keefe, J. (2006). The Hippocampus Book. (Oxford University Press).

[6] Arnatkevi, A. (2018). A practical guide to linking brain-wide gene expression and neuroimaging data Keywords Measuring gene expression.

[7] Ashburner, J. (2007). A fast diffeomorphic image registration algorithm. NeuroImage, 38, 95–113.

[8] Bienkowski, M. S., Bowman, I., Song, M. Y., Gou, L., Ard, T., Cotter, K., Zhu, M., Benavidez, N. L., Yamashita, S., Abu-Jaber, J., Azam, S., Lo, D., Foster, N. N., Hintiryan, H., & Dong, H.-W. (2018). Integration of gene expression and brain-wide connectivity reveals the multiscale organization of mouse hippocampal networks. Nature Neuroscience.

[9] Breunig, J. J., Sarkisian, M. R., Arellano, J. I., Morozov, Y. M., Ayoub, A. E., Sojitra, S., Wang, B., Flavell, R. A., Rakic, P., & Town, T. (2008). Primary cilia regulate hippocampal neurogenesis by mediating sonic hedgehog signaling. Proceedings of the National Academy of Sciences, 105, 13127–13132.

[10] Brunec, I. K., Bellana, B., Ozubko, J. D., Man, V., Robin, J., Liu, Z.-x., Grady, C., Rosenbaum, R. S., Winocur, G., Barense, M. D., & Moscovitch, M. (2018). Multiple Scales of Representation along the Hippocampal Anteroposterior Axis in Humans. Current Biology, 28, 2129–2135.

[11] Buckner, R. L. & Krienen, F. M. (2013). The evolution of distributed association networks in the human brain. Trends in Cognitive Sciences, 17, 648–665.

[12] Buxbaum, J. N., Ye, Z., Reixach, N., Friske, L., Levy, C., Das, P., Golde, T., Masliah, E., Roberts, A. R., & Bartfai, T. (2008). Transthyretin protects Alzheimer’s mice from the behavioral and biochemical effects of A toxicity. Proceedings of the National Academy of Sciences, 105, 2681–2686.

[13] Cembrowski, M. S., Bachman, J. L., Wang, L., Sugino, K., Shields, B. C., Cembrowski, M. S., Bachman, J. L., Wang, L., Sugino, K., Shields, B. C., & Spruston, N. (2016). Spatial Gene-Expression Gradients Underlie Prominent Heterogeneity of CA1 Pyramidal Neurons Article Spatial Gene-Expression Gradients Underlie Prominent Heterogeneity of CA1 Pyramidal Neurons. Neuron, 89, 351–368.

[14] Chase, H. W., Clos, M., Dibble, S., Fox, P., Grace, A. A., Phillips, M. L., & Eickhoff, S. B. (2015). Evidence for an anterior-posterior differentiation in the human hippocampal formation revealed by meta-analytic parcellation of fMRI coordinate maps: Focus on the subiculum. NeuroImage, 113, 44–60.

[15] Christensen, T., Bisgaard, C. F., Nielsen, H. B., & Wiborg, O. (2010). TRANSCRIPTOME DIFFERENTIATION ALONG THE DORSO VENTRAL AXIS IN LASER-CAPTURED MICRODIS-SECTED RAT HIPPOCAMPAL GRANULAR CELL LAYER. NSC, 170, 731–741.

[16] Collin, S. H. P., Milivojevic, B., & Doeller, C. F. (2015). Memory hierarchies map onto the hippocampal long axis in humans. Nature Neuroscience, 18, 1562–1564.

[17] Cuenco, K. T., Friedland, R., Baldwin, C. T., Guo, J., Vardarajan, B., Lunetta, K. L., Cupples, L. A., Green, R. C., DeCarli, C., & Farrer Lindsay A., L. A. (2011). Association of TTR polymorphisms with hippocampal atrophy in Alzheimer disease families. Neurobiology of Aging, 32, 249–256.

[18] Diamandis, E. P., Scorilas, A., Kishi, T., Blennow, K., Luo, L. Y., Soosaipillai, A., Rademaker, A. W., & Sjogren, M. (2004). Altered kallikrein 7 and 10 concentrations in cerebrospinal fluid of patients with Alzheimer’s disease and frontotemporal dementia. Clinical Biochemistry, 37, 230–237.

[19] Dong, H.-w., Swanson, L. W., Chen, L., Fanselow, M. S., & Toga, A. W. (2009). Genomic anatomic evidence for distinct functional domains in hippocampal field CA1. 106.

[20] Fanselow, M. S. & Dong, H.-w. (2010). Review Are the Dorsal and Ventral Hippocampus Functionally Distinct Structures? Neuron, 65, 7–19.

[21] Fornito, A., Arnatkevčiūtė, A., & Fulcher, B. D. (2018). Bridging the Gap between Connectome and Transcriptome. Trends in Cognitive Sciences, xx, 1–17.

[22] Fuentealba, P., Klausberger, T., Karayannis, T., Suen, W. Y., Huck, J., Tomioka, R., Rockland, K., Capogna, M., Studer, M., Morales, M., & Somogyi, P. (2010). Expression of COUP-TFII Nuclear Receptor in Restricted GABAergic Neuronal Populations in the Adult Rat Hippocampus. Journal of Neuroscience, 30, 1595–1609.

[23] Giaccio, R. G. (2006). The dual origin hypothesis: An evolutionary brain-behavior framework for analyzing psychiatric disorders. 30, 526–550.

[24] Hastie, T., Tibshirani, R., & Wainwright, M. (2015). Statistical learning with sparsity: the lasso and generalization. (CRC Press).

[25] Haufe, S., Meinecke, F., Görgen, K., Dähne, S., Haynes, J. D., Blankertz, B., & Bießmann, F. (2014). On the interpretation of weight vectors of linear models in multivariate neuroimaging. NeuroImage, 87, 96–110.

[26] Hawrylycz, M., Miller, J. A., Menon, V., Feng, D., Dolbeare, T., Guillozet-Bongaarts, A. L., Jegga, A. G., Aronow, B. J., Lee, C.-K., Bernard, A., Glasser, M. F., Dierker, D. L., Menche, J., Szafer, A., Collman, F., Grange, P., Berman, K. A., Mihalas, S., Yao, Z., Stewart, L., Barabási, A.-L., Schulkin, J., Phillips, J., Ng, L., Dang, C., Haynor, D. R., Jones, A., Van Essen, D. C., Koch, C., & Lein, E. (2015). Canonical genetic signatures of the adult human brain. Nature neuroscience, 18, 1832–1844.

[27] Hu, J. S., Vogt, D., Lindtner, S., Sandberg, M., Silberberg, S. N., & Rubenstein, J. L. R. (2017). ¡i¿Coup-TF1¡/i¿ and ¡i¿Coup-TF2¡/i¿ control subtype and laminar identity of MGE-derived neocortical interneurons. Development, 144, 2837–2851.

[28] LaJoie, R., Landeau, B., Perrotin, A., Bejanin, A., Egret, S., P??lerin, A., M??zenge, F., Belliard, S., deLaSayette, V., Eustache, F., Desgranges, B., & Ch??telat, G. (2014). Intrinsic connectivity identifies the hippocampus as a main crossroad between alzheimer’s and semantic dementia-targeted networks. Neuron, 81, 1417–1428.

[29] Lee, A.-r., Kim, J.-h., Cho, E., Kim, M., Park, M., & Albrecht, A. (2017). Dorsal and Ventral Hippocampus Differentiate in Functional Pathways and Differentially Associate with Neurological Disease-Related Genes during Postnatal Development. 10, 1–14.

[30] Lee, H., Park, Y. M., & Lee, S. (2015). Principal Component Regression by Principal Component Selection. Communications for Statistical Applications and Methods, 22, 173–180.

[31] Leonardo, E., Richardson-Jones, J., Sibille, E., Kottman, A., & Hen, R. (2006). Molecular heterogeneity along the dorsalventral axis of the murine hippocampal CA1 field: a microarray analysis of gene expression. Neuroscience, 137, 177–186.

[32] Lin, F. J., Qin, J., Tang, K., Tsai, S. Y., & Tsai, M. J. (2011). Coup d’Etat: An orphan takes control. Endocrine Reviews, 32, 404–421.

[33] Lladó, A., Tort-merino, A., Sánchez-valle, R., Falgàs, N., Balasa, M., Bosch, B., Castellví, M., Olives, J., Antonell, A., & Hornberger, M. (2018). Neurobiology of Aging The hippocampal longitudinal axis d relevance for underlying tau and TDP-43 pathology. Neurobiology of Aging, 70, 1–9.

[34] Lourenço, F. C., Galvan, V., Fombonne, J., Corset, V., Llambi, F., Müller, U., Bredesen, D. E., & Mehlen, P. (2009). Netrin-1 interacts with amyloid precursor protein and regulates amyloid-*β* production. Cell Death and Differentiation, 16, 655–663.

[35] Marcus, D. S., Wang, T. H., Parker, J., Csernansky, J. G., Morris, J. C., & Buckner, R. L. (2007). Open Access Series of Imaging Studies (OASIS): Cross-sectional MRI Data in Young, Middle Aged, Nondemented, and Demented Older Adults. Journal of Cognitive Neuroscience, 19, 1498–1507.

[36] Margulies, D. S., Ghosh, S. S., Goulas, A., Falkiewicz, M., & Huntenburg, J. M. (2016). Situating the default-mode network along a principal gradient of macroscale cortical organization. 113, 12574–12579.

[37] McKhann, G., Knopman, D. S., Chertkow, H., Hymann, B., Jack, C. R., Kawas, C., Klunk, W., Koroshetz, W., Manly, J., Mayeux, R., Mohs, R., Morris, J., Rossor, M., Scheltens, P., Carrillo, M., Weintrub, S., & Phelphs, C. (2011). The diagnosis of dementia due to Alzheimers disease: Recommendations from the National Institute on Aging-Alzheimers Association workgroups on diagnostic guidelines for Alzheimers disease. Alzheimers Dementia, 7, 263–269.

[38] Neary, D., Snowden, J. S., Gustafson, L., Passant, U., Stuss, D., Black, S., Freedman, M., Kertesz, A., Robert, P. H., Albert, M., Boone, K., Miller, B. L., Cummings, J., & Benson, D. F. (1998). Frontotemporal lobar degeneration: a consensus on clinical diagnostic criteria. Neurology, 51, 1546–54.

[39] Nordin, K., Persson, J., Stening, E., Herlitz, A., Larsson, E. M., & Söderlund, H. (2018). Structural whole-brain covariance of the anterior and posterior hippocampus: Associations with age and memory. Hippocampus, 28, 151–163.

[40] Ossenkoppele, R., Schonhaut, D. R., Schöll, M., Lockhart, S. N., Ayakta, N., Baker, S. L., O’Neil, J. P., Janabi, M., Lazaris, A., Cantwell, A., Vogel, J., Santos, M., Miller, Z. A., Bettcher, B. M., Vossel, K. A., Kramer, J. H., Gorno-Tempini, M. L., Miller, B. L., Jagust, W. J., & Rabinovici, G. D. (2016). Tau PET patterns mirror clinical and neuroanatomical variability in Alzheimer’s disease. Brain, 139, 1551–1567.

[41] Poldrack, R. A., Mumford, J. A., Schonberg, T., Kalar, D., & Barman, B. (2012). Discovering Relations Between Mind, Brain, and Mental Disorders Using Topic Mapping. 8.

[42] Poppenk, J., Evensmoen, H. R., Moscovitch, M., & Nadel, L. (2013). Long-axis specialization of the human hippocampus. Trends in Cognitive Sciences, 17, 230–240.

[43] Rama, N., Goldschneider, D., Corset, V., Lambert, J., Pays, L., & Mehlen, P. (2012). Amyloid precursor protein regulates netrin-1-mediated commissural axon outgrowth. Journal of Biological Chemistry, 287, 30014–30023.

[44] Ranganath, C. & Ritchey, M. (2012). Two cortical systems for memory-guided behaviour. Nature Reviews Neuroscience, 13, 713–726.

[45] Reardon, P. K., Seidlitz, J., Vandekar, S., Liu, S., Patel, R., Park, M. T., Alexander-Bloch, A., Clasen, L. S., Blumenthal, J. D., Lalonde, F. M., Giedd, J. N., Gur, R., Gur, R., Lerch, J. P., Chakravarty, M. M., Satterthwaite, T., Shinohara, R. T., & Raznahan, A. (2018). Normative brain size variation and brain shape diversity in humans.

[46] Reilly, K. C. O., Flatberg, A., & Islam, S. (2015). Identification of dorsal ventral hippocampal differentiation in neonatal rats. Brain Structure and Function, pp. 2873–2893.

[47] Richiardi, J., Altmann, A., Milazzo, A.-C., Chang, C., Chakravarty, M. M., Banaschewski, T., Barker, G. J., Bokde, A. L. W., Bromberg, U., Büchel, C., Conrod, P., Fauth-Bühler, M., Flor, H., Frouin, V., Gallinat, J., Garavan, H., Gowland, P., Heinz, A., Lemaître, H., Mann, K. F., Martinot, J.-L., Nees, F., Paus, T., Pausova, Z., Rietschel, M., Robbins, T. W., Smolka, M. N., Spanagel, R., Ströhle, A., Schumann, G., Hawrylycz, M., Poline, J.-B., Greicius, M. D., & consortium, I. (2015). BRAIN NETWORKS. Correlated gene expression supports synchronous activity in brain networks. Science (New York, N.Y.), 348, 1241–1244.

[48] Romero-garcia, R., Whitaker, K. J., Seidlitz, J., Shinn, M., Fonagy, P., Dolan, R. J., Jones, P. B., Goodyer, I. M., Consortium, N., Bullmore, E. T., & Petra, E. V. (2018). NeuroImage Structural covariance networks are coupled to expression of genes enriched in supragranular layers of the human cortex. 171, 256–267.

[49] Strange, B. A., Witter, M. P., Lein, E. S., & Moser, E. I. (2014). Functional organization of the hippocampal longitudinal axis. Nature Reviews Neuroscience, 15, 655–669.

[50] Sunkin, S. M., Ng, L., Lau, C., Dolbeare, T., Gilbert, T. L., Thompson, C. L., Hawrylycz, M., & Dang, C. (2012). Allen Brain Atlas: an integrated spatio-temporal portal for exploring the central nervous system. Nucleic Acids Research, 41, D996–D1008.

[51] Thompson, C. L., Pathak, S. D., Jeromin, A., Ng, L. L., Macpherson, C. R., Mortrud, M. T., Cusick, A., Riley, Z. L., Sunkin, S. M., Bernard, A., Puchalski, R. B., Gage, F. H., Jones, A. R., Bajic, V. B., Hawrylycz, M. J., & Lein, E. S. (2008). Article Genomic Anatomy of the Hippocampus. Neuron, 60, 1010–1021.

[52] Vos de Wael, R., Larivìere, S., Caldairou, B., Hong, S.-J., Margulies, D. S., Jefferies, E., Bernasconi, A., Smallwood, J., Bernasconi, N., & Bernhardt, B. C. (2018). Anatomical and microstructural determinants of hippocampal subfield functional connectome embedding. Proceedings of the National Academy of Sciences, 115, 201803667.

[53] Wager, T. D., Atlas, L. Y., Lindquist, M. A., Roy, M., Woo, C.-W., & Kross, E. (2013). An fMRI-Based Neurologic Signature of Physical Pain. New England Journal of Medicine, 368, 1388–1397.

[54] Wang, G. Z., Belgard, T. G., Mao, D., Chen, L., Berto, S., Preuss, T. M., Lu, H., Geschwind, D. H., & Konopka, G. (2015). Correspondence between Resting-State Activity and Brain Gene Expression. Neuron, 88, 659–666.

[55] Yeo, B., Krienen, F., Sepulcre, J., Sabuncu, M., Lashkari, D., Hollinshead, M., Roffman, J., Smoller, J., Zollei, L., Polimeni, J., Fischl, B., Liu, H., & Buckner, R. (2011). The organization of the human cerebral cortex estimated by intrinsic functional connectivity. Journal of Neuroph, 106, 1125–1165.

[56] Zeisel, A., Munoz-Manchado, A. B., Codeluppi, S., Lonnerberg, P., La Manno, G., Jureus, A., Marques, S., Munguba, H., He, L., Betsholtz, C., Rolny, C., Castelo-Branco, G., Hjerling-Leffler, J., & Linnarsson, S. (2015). Cell types in the mouse cortex and hippocampus revealed by single-cell RNA-seq. Science, 347, 1138–1142.

